# Discovering patterns of pleiotropy in genome-wide association studies

**DOI:** 10.1101/273540

**Authors:** Jianan Zhana, CHARGE ECG Working Group, Jessica van Setten, Jennifer Brody, Brenton Swenson, Anne M. Butler, Harry Campbell, Fabiola Del Greco, Daniel S. Evans, Quince Gibson, Daniel F. Gudbjartsson, Kathleen F. Kerr, Bouwe P. Krijthe, Leo-Pekka Lyytikäinen, Christian Müller, Martina Müller-Nurasyid, Ilja M. Nolte, Sandosh Padmanabhan, Marylyn D. Ritchie, Antonietta Robino, Albert V. Smith, Maristella Steri, Toshiko Tanaka, Alexander Teumer, Stella Trompet, Sheila Ulivi, Niek Verweij, Xiaoyan Yin, David O. Arnar, Folkert W. Asselbergs, John Barnard, Josh Bis, Stefan Blankenberg, Eric Boerwinkle, Yuki Bradford, Brendan M. Buckley, Mina K. Chung, Dana Crawford, Marcel den Hoed, Josh Denny, Anna F. Dominiczak, Georg B. Ehret, Mark Eijgelsheim, Patrick Ellinor, Stephan B. Felix, Lude Franke, Tamara B. Harris, Susan R. Heckbert, Hilma Holm, Unnur Thorsteinsdottir, Gandin Ilaria, Annamaria Iorio, Mika Kähönen, Ivana Kolcic, Jan A. Kors, Edward G. Lakatta, Lenore J. Launer, Honghuang Lin, Henri J. Lin, Yongmei Liu, Ruth Loos, Steve Lubitz, Peter MacFarlane, Jared W. Magnani, Irene Mateo Leach, Thomas Meitinger, Braxton Mitchell, Thomas Munzel, George J. Papanicolaou, Annette Peters, Arne Pfeufer, Peter M. Pramstaller, Olli T. Raitakari, Jerome I. Rotter, Igor Rudan, Nilesh J. Samani, David Schlessinger, Claudia T. Silva Aldana, Moritz Sinner, Jonathan D. Smith, Harold Snieder, Elsayed Soliman, Timothy D. Spector, David J. Stott, Konstantin Strauch, Kirill V. Tarasov, Andre G. Uitterlinden, David R. van Wagoner, Uwe Völker, Henry Völzke, Melanie Waldenberger, Harm Jan Westra, Philipp S. Wild, Tanja Zeller, Alvaro Alonso, Christy L. Avery, Stefania Bandinelli, Emelia J. Benjamin, Francesco Cucca, Steven R. Cummings, Marcus Dörr, Luigi Ferrucci, Paolo Gasparini, Vilmundur Gudnason, Carolina Hayward, Andrew A. Hicks, Yalda Jamshidi, J. Wouter Jukema, Stefan Kääb, Terho Lehtimäki, Patricia B. Munroe, Afshin Parsa, Ozren Polasekd, Bruce Psaty, Dan Roden, Renate B. Schnabel, Gianfranco Sinagra, Kari Stefansson, Bruno H. Stricker, Pim van der Harst, Cornelia M. van Duijn, James F. Wilson, Sina Gharib, Paul I.W. de Bakker, Aaron Isaacs, Dan E. Arking, Nona Sotoodehnia, Dan E. Arking, Joel S. Baderab

**Author notes:** Corresponding author Email address (Joel S. Bader).

## Abstract

**Motivation:** Genome-wide association studies have had great success in identifying human genetic variants associated with disease, disease risk factors, and other biomedical phenotypes. Many variants are associated with multiple traits, even after correction for trait-trait correlation. Discovering subsets of variants associated with a shared subset of phenotypes could help reveal disease mechanisms, suggest new therapeutic options, and increase the power to detect additional variants with similar pattern of associations. Here we introduce two methods based on a Bayesian framework, SNP And Pleiotropic PHenotype Organization (SAPPHO), one modeling independent phenotypes (SAPPHO-I) and the other incorporating a full phenotype covariance structure (SAPPHO-C). These two methods learn patterns of pleiotropy from genotype and phenotype data, using identified associations to discover additional associations with shared patterns.

**Results:** The SAPPHO methods, along with other recent approaches for pleiotropic association tests, were assessed using data from the Atherosclerotic Risk in Communities (ARIC) study of 8,000 individuals, whose gold-standard associations were provided by meta-analysis of 40,000 to 100,000 individuals from the CHARGE consortium. Using power to detect gold-standard associations at genome-wide significance (0.05 family-wise error rate) as a metric, SAPPHO performed best. The SAPPHO methods were also uniquely able to select the most significant variants in a parsimonious model, excluding other less likely variants within a linkage disequilibrium block. For meta-analysis, the SAPPHO methods implement summary modes that use sufficient statistics rather than full phenotype and genotype data. Meta-analysis applied to CHARGE detected 16 additional associations to the gold-standard loci, as well as 124 novel loci, at 0.05 false discovery rate. Reasons for the superior performance were explored by performing simulations over a range of scenarios describing different genetic architectures. With SAPPHO we were able to learn genetic structures that were hidden using the traditional univariate tests.

**Availability:** https://bitbucket.org/baderlab/fast/wiki/Home. SAPPHO software is available under the GNU General Public License, v2.

## 1. Introduction

Genome-wide association studies (GWAS) have had remarkable success in identifying genetic variants responsible for human disease, disease risk factors, and other biomedical phenotypes. To date, more than 17607 variants, primarily single nucleotide polymorphisms (SNPs), have been associated at genome-wide significance with at least 785 distinct traits, according to the GWAS catalog [1]. Many variants are pleiotropic, with significant associations with multiple traits (Fig. 1). Observations of pleiotropy motivate systematic approaches to identify pleiotropic variants. Such approaches could use observed patterns of pleiotropy to identify additional variants that follow the same pattern. In a Bayesian setting, the observed patterns would provide prior probabilities that could boost the confidence that other variants with the same pattern are true associations, even if their univariate p-values do not reach conventional genomewide significance thresholds. A second valuable application could be to use pleiotropic associations to infer mechanisms shared by multiple diseases, which could lead to new therapeutic approaches including drug repurposing.

**Figure 1:**
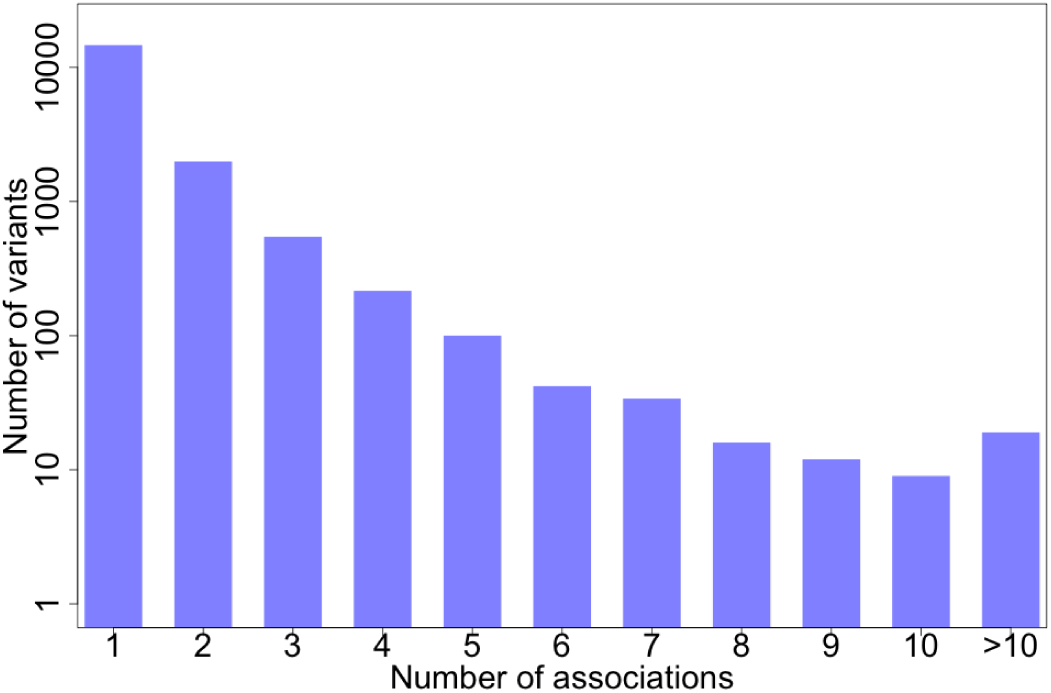
Histogram of pleiotropic variants. Counts are for variants associated with at least one trait at genome-wide significance (*p* < 5 × 10^−8^) [1]; no attempt was made to correct for correlation between variants (linkage disequilibrium) or between traits. The number of total variants is 17607, and total number of phenotypes is 785.

A challenge is that we do not know in general which traits share causal genetic factors. While these pleiotropic patterns may be discovered from genomewide association study data, re-use of the same data for pattern discovery and association discovery requires new methods to control false discovery rates. A second important challenge is to develop methods that provide as direct a route as possible to the most significant variant within an association locus. Methods that produce parsimonious models, selecting just the most significant variant and excluding the neighboring linked variants, have great value. A third challenge is to incorporate the phenotype-phenotype covariance structure in the analysis to discriminate between a model in which a variant affects two phenotypes directly and an alternative model in which a variant directly affects one phenotype, which in turn affects a second correlated phenotype.

While there are no pleiotropic association methods in general use, there have been three general directions for methods development. First, for small collections of highly correlated phenotypes, a reasonable approach is to aggregate associations over all the phenotypes. This approach has greatest power when the true model is that a variant affects each phenotype. A recent report demonstrated good power for phenotypes related to hypertension [2]. This approach is similar in motivation to gene-based methods for GWAS signal aggregation, including VEGAS for common variants [3] and SKAT for rare variants [4].

When phenotypes are highly correlated and possibly redundant, a second direction has been to use orthogonalization methods, usually principal component analysis or singular value decomposition, to identify a rank-reduced set of linear combinations. A representative method is principal components of heritability (PCH), which generates linear combination of phenotypes with highest heritability for each genetic variant [5]. A drawback of this approach is that validating an association with a linear combination of phenotypes is more difficult than validating an association with a single phenotype, particularly when different studies assess different sets of phenotypes. An alternative approach is to use a linear combination of phenotypes as a feature to predict the variant genotype, reversing the typical direction of regression. The MultiPhen method uses ordinal regression to perform this type of test, with increased power for variants affecting multiple phenotypes [6]. While methods such as canonical correlation analysis (CCA) and MANOVA that assume that genotypes follow a normal distribution have inflated type-I error, MultiPhen produces no such inflation when parametric p-values are used.

A third approach has been to adapt methods like L1/LASSO [7] that favor selection of sparser sets of features [8]. Despite the promise of this approach, it has not been widely used, possibly because the L1 regularization is still too weak to reject variants introduced through correlation only and not causation and because of computational costs for genome-wide applications.

The approach pursued here is to exploit observed association patterns to identify additional variants following the same pattern. These patterns are biclusters, with subsets of variants associated with subsets of phenotypes. Biclustering has been a productive approach for identifying block structure in gene expression data [9]. Biclustering is not directly applicable to GWAS data, however, because blocks of SNPs identified by näive application of standard biclustering algorithms would be dominated by non-causal variants in linkage disequilibrium or haplotype blocks with a single causal variant.

We report results for a new Bayesian framework for genome-wide association studies of multiple phenotypes with shared genetics: SNP And Pleiotropic PHenotype Organization, SAPPHO. The SAPPHO method is motivated by our previous work developing a Bayesian method for gene-based association tests, Gene-Wide Significance (GWiS) [10, 11], which aggregates statistical associations of multiple independent variants within a gene for a single phenotype. Each identified variant within a gene effectively updates the prior probability that additional variants within the same gene are also associated, permitting successive identification of variants with smaller effects that could be missed by conventional univariate tests of individual SNPs. Using an assessment with real data and gold standards from meta-analysis, GWiS was found to have greater power than univariate tests and also greater power than other gene-based methods, including methods based on summing the effects over all variants [3] and methods using L1/LASSO [7]. GWiS had robust performance across different genetic architectures, including the number of true effects per gene and the minor allele frequencies. The robust performance was in part due to a lack of tuning parameters. Instead, most parameters in the GWiS model were treated as nuisance parameters and removed by integration.

The SAPPHO method uses a similar approach to associate individual variants with multiple phenotypes in a single genome-wide model for testing *T* total SNPs for association with *P* total phenotypes. Model priors interpolate between two probability distributions for genetic architecture, one which each of the *T* × *P* possible SNP-phenotype associations is independent, and a second in which each of the 2^*P*^ possible patterns of association has its own prior probability. All of the remaining association structure parameters are integrated out, yielding a method with only a single adjustable parameter, the mixing fraction of the two priors. This parameter is essentially the threshold for the weakest possible variant-phenotype association that can be entered into a regression model. Identifying the most likely model is NP-hard, and heuristics are needed for an acceptable runtime. SAPPHO uses a greedy forward approach to identify a local optimum with an algorithm that scales linearly with the number of SNP-phenotype associations identified in the data.

We evaluate our proposed method through analysis of cardiovascular electrocardiogram (ECG) phenotypes and simulation. For ECG phenotypes, metaanalyses of studies with 40,000 to 100,000 individuals have been conducted with overlapping sets of variants associated with PR, QRS, and QT intervals. Notwithstanding concerns about missing heritability [12], the fraction of heritability explained by genome-wide significant associations for these traits ranges from 4% to 17% [13, 14, 15, 16]. The genome-wide significant findings provided by meta-analysis provide gold-standard true positive associations for assessing the power of different methods. The SAPPHO methods have the further potential to provide new biomedical knowledge by revealing classes of variants that contribute to distinct subsets of ECG parameters.

The Methods section provides a mathematical description of SAPPHO and the algorithm used to identify an optimum in the space of all possible models. Briefer summaries of other approaches are provided, together with summaries of real and simulated data used for assessment. The Results section reports on the assessment results and the pleiotropic patterns observed for ECG traits and also for simulations representing a range of scenarios of phenotypes that share genetic and environmental factors. The Discussion concludes with an interpretation of the benefits and drawbacks of different pleiotropy methods and a vision for possible future directions to discover and exploit pleiotropy in human genetic association studies.

## Methods

### 2.1 Genetic model

SAPPHO has been developed for quantitative phenotypes. Case/control or other dichotomous phenotypes can be represented as 0/1 encodings, which general retain high power when causal variants have small effects [11]. Similarly, rank-ordered categories can be represented as corresponding integers. Unranked categories can in principle be represented as 0/1 indicators for each category; in practice, these are less common than quantitative, graded, or dichotomous phenotypes. Extensions to dichotomous phenotypes or general linear models in the exponential family are possible but more computationally intensive without a corresponding gain in power [11].

The SAPPHO statistical model considers a population of *N* unrelated individuals and *P* distinct phenotypes, with each individual assessed for each phenotype. The phenotype data is represented as a real-valued phenotype matrix **Y** with *N* rows and *P* columns. Individuals are also genotyped at distinct loci corresponding to *T* total independent tests, represented as a real-valued genotype matrix **X** with *N* rows and *T* columns. In most applications, the genotype values will correspond to allele frequencies or dosages for bi-allelic single nucleotide polymorphisms (SNPs), measured directly or imputed. All data elements of **Y** and **X** are assumed present. In practice, some individuals will lack data for some genotypes and phenotypes. In this work, for simplicity only individuals with complete data are retained. Exclusion could be done at the level of individual means, variances, and covariances of phenotypes and genotypes, which in theory leads to non-positive-definite covariance matrices but in practice usually does not cause numerical instabilities [11].

An association model, denoted *M*, specifies the direct effects of variants on phenotypes. For this work, we restrict attention to linear models. Thus, a model specifies which elements of a regression coefficient matrix ***β*** with *T* rows and *P* columns may be non-zero; the number of non-zero elements is denoted *|M |*. The model does not specify the corresponding values; these are treated as nuisance parameters that are integrated out. We consider two different models representing alternative assumptions about the phenotype covariance matrix: SAPPHO-I models each phenotype as independent given the genetic effects; SAPPHO-C models the complete phenotype-phenotype covariance structure. Given the model *M* and the genotypes **X** for an individual, the probability distribution for phenotypes **Y** is multivariate normal with covariance matrix **Ω**, diagonal for SAPPHO-I and including off-diagonal elements for SAPPHO-C,

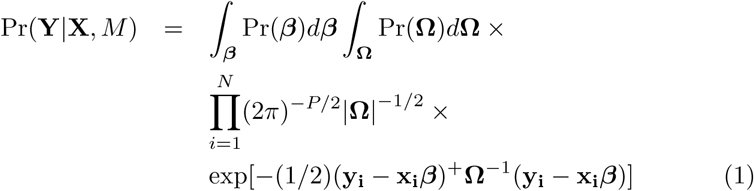

where **x_i_** and **y_i_** are genotype and phenotype vectors for individual *i*, and the superscript ^+^ denotes transpose. The integral over ***β*** is over the *|M |* nonzero elements, and the ***β*** and **Ω** integrals include formal normalization factors Pr(***β***) and Pr(**Ω**). The notation *|***Ω***|* denotes the determinant of the phenotype covariance matrix **Ω**. This covariance matrix does not include the genetic contributions; the observed covariance matrix **V** is

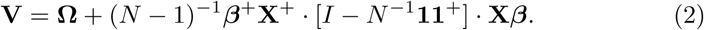

The normalization factors Pr(***β***) and Pr(**Ω**) formally depend on meta-parameters for regularization. In practice, we use the asymptotic limit that excludes the contribution of the meta-parameters, as we did with GWiS [10], keeping terms of order ln *N* and greater. Performing steepest descents around the maximum likelihood estimates 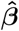 and 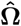, equivalent to the Bayesian Information Criterion or BIC [18], the asymptotic limit for the log-probability of the observed phenotypes given genotypes and model is

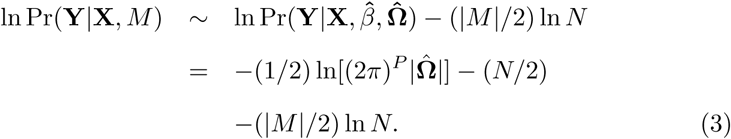

For SAPPHO-I, we assume that the phenotypes are independent with each other; this is essentially equivalent to setting the non-diagonal elements of **Ω** equal to zero, leaving only the non-diagonal 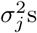 that correspond to the residual variance of each phenotype; in other words, now 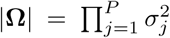, where 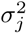 is the residual variance of each phenotype. The probability distribution for phenotypes **Y** becomes a product of *P* normal distributions, with the number of distributions equal to the number of phenotypes:

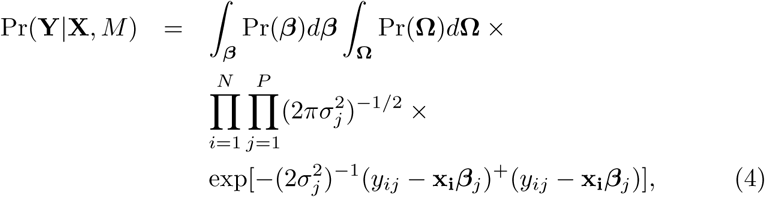

where *y*_*ij*_ is the *j*th element of vector **y_i_**, and ***β***_*j*_ denotes the *j*th column of matrix ***β***. Adopting the same asymptotic approximation for the log-probability using BIC yields

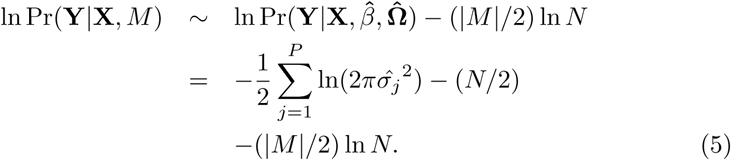

### 2.2. Model prior

The model prior probability, Pr(*M*), is represented as the product of two terms,

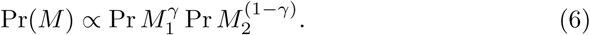

This form corresponds to linear interpolation on a logarithmic scale,

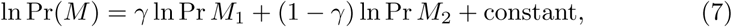

where *γ ∈* [0, 1] and the constant term is model-independent and may be ignored. The prior Pr *M*_1_ penalizes each genotype-phenotype association individually, while the prior Pr *M*_2_ penalizes based on each different association pattern, as described below.

The prior Pr *M*_1_ models each possible SNP-phenotype pair as a binary random variable reflecting association with probability *θ* or no association with probability 1 – *θ* for *θ ∈* [0, 1]; the single parameter *θ* is shared by each of the *T* × *P* possible genotype-phenotype associations. For any model with *|M |* = *K* total associations, Pr *M*_1_ is

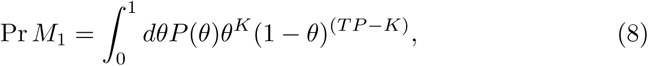

where *P* (*θ*) is a possible prior on *θ*; we use the uniform prior *P* (*θ*) = 1. While *P* varies with the number of phenotypes in different studies, *T* is set equal to the conventional number of independent effects in the genome for human GWAS, 10^6^. The nuisance parameter *θ* is integrated out to yield the standard result,

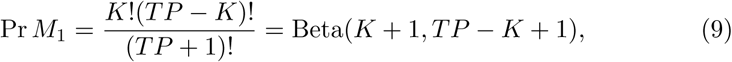

where ‘!’ denotes the factorial function and ‘Beta’ is the standard Beta function extending the combinatorial factor to non-integer arguments.

The derivation of Pr *M*_2_ is similar to Pr *M*_1_, except that it considers patterns rather than individual associations. A pattern *α* is one of the 2^*P*^ *–* 1 possible subsets of phenotypes, excluding the null pattern of no associations. The probability that a SNP belongs to pattern *α* is denoted *θ*_*α*_, with *θ*_*α*_ *∈* [0, 1] and Σ_*α*_ *θ*_*α*_ = 1 defining a multinomial probability distribution. Denoting *n*_*α*_ as the total number of variants with pattern *α*, and Σ_*α*_ *n*_*α*_ = *n*, *n* being the total number of associated SNPs, the probability for a particular model *M* is

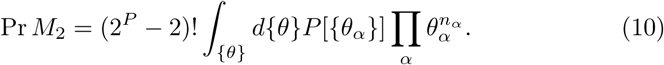

The integral is over all possible feasible parameters and *P* [*{θ*_*α*_*}*] is a possible prior distribution; we use the uniform distribution *P* [*{θ*_*α*_*}*] = 1. The term (2^*P*^ – 2)! is the standard normalization factor for a multinomial distribution. The nuisance parameters *θ*_*α*_ are removed by integration, yielding the standard result,

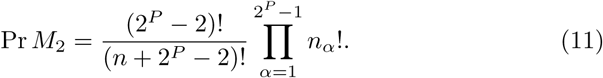

Note that for Pr *M*_2_ only the occupied patterns contribute to the model probability, similar to latent Dirichlet allocation (LDA) in which only occupied states contribute [19]. This probability model favors pleiotropy models in which variants share the same association patterns. The overall prefactor (2^*P*^ – 2)!/(*n* + 2^*P*^ – 2)! is identical for all models and independent of the occupation numbers *{n*_*α*_*}* for different patterns. Therefore, for the purpose of computational efficiency, we use

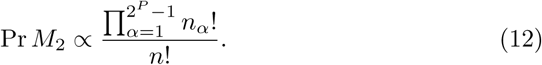

### 2.3. Model score

The goal of SAPPHO is to identify the most likely model, 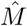, defined as 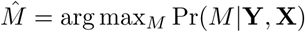. The posterior probability of a model is defined by Bayes rule as

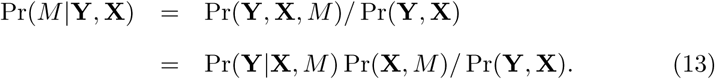

We make the standard assumption that the model *M* is independent of the genotype data, Pr(**X***, M*) = Pr(**X**) Pr(*M*), giving

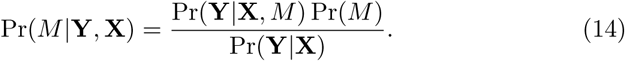

The conditional probability Pr(**Y***|***X**) is independent of model and need not be calculated. Similarly, to avoid numeric overflow and underflow, model posterior probabilities are always calculated as log-likelihood ratios relative to the null model, defined as the model score *S*_*M*_,

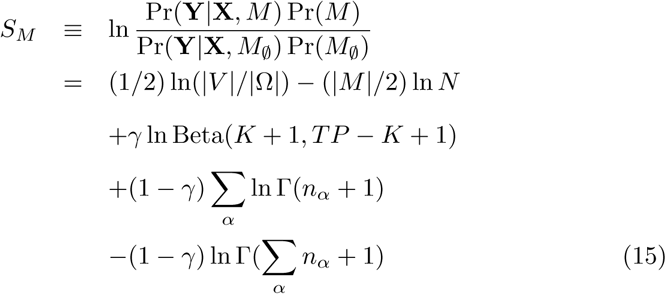

In practice, all models, including the null model, typically include constant terms for phenotype mean and covariates that represent relevant clinical variables, including sex, age, height, weight, and body mass index, and possible additional covariates that describe population structures. Regression coefficients for these covariates are calculated in parallel with regression coefficients for genetic variants, but they make no net contribution when models are compared. For computational efficiency, SAPPHO regresses out the known covariates and then operates on the residuals.

The parameter *γ* is the single adjustable parameter in the SAPPHO method. While it could be set using cross-validation, this would require a gold-positive training set and depends on the genetic architecture. An architecture with no shared genetic factors would favor Pr *M*_1_, whereas an architecture with a small number of strong patterns would favor Pr *M*_2_. Instead, we relate *γ* to the effect size required to enter a new SNP-phenotype association into a model. To be more specific, we take the dominant term from the Beta penalty, together with the BIC penalty, giving the *χ*^2^ threshold for adding a single association to the model,

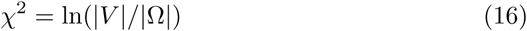

The value of *γ* is then calculated accordingly,

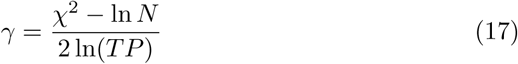

Behavior of SAPPHO under different values of this tuning parameter is discussed in the following paragraphs. In general, with real data, we found that setting *γ* ln(*TP*) = ln(10^4^) is a good value to control for type I and II error rate; with simulation, different tuning parameter will lead to different behaviors of SAPPHO, favoring different true underlying real association patterns.

### 2.4. Model search and variant ranking

Identifying 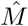 is NP-hard and is not attempted directly. Instead, a greedy forward approach is employed. Given a current model, all models that may be reached by adding a single genetic association are considered. There are two possible cases. In one case, a variant with no associations gains an association to a single phenotype. In the second case, a variant associated with a subset of phenotypes gains an association with one additional phenotype. With *T* total variants and *P* total phenotypes, this procedure requires calculating the posterior probability for approximately *T* × *P* possible models. The model with the greatest increase in posterior probability is selected, and the procedure continues until any additional association decreases the posterior probability. The resulting model, locally optimal with respect to adding associations, is termed 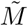, distinct from the global optimum 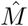. In principle, stepwise forwardbackward selection would also be possible, but would require full matrix inverses that would vastly increase the computational cost. Backward steps removing a variant and all of its associations are used at the very end of the model search, however, for variant ranking.

For SAPPHO-I, computation is made more efficient by using successive orthogonalization rather than matrix inverses at each step to obtain the new maximum likelihood estimates for 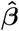 and 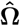.

To further speed the model search by SAPPHO-I and SAPPHO-C, the candidate list of variants is pre-filtered, as is common with with other approaches that build multivariate genome-scale models. A preliminary univariate test is conducted for each of the *T* total variants with each of the *P* total phenotypes, yielding *T* × *P* total p-values. Each variant is then assigned its minimum p-value across the *P* phenotypes, and variants are sorted from smallest to largest p-value. Candidate variants are taken from this list in increasing p-value order, with variants excluded if they are in strong linkage disequilibrium (*r*^2^ ≥ 0.8) with a better-ranked variant that has already been selected. Selection ends when the p-value of the best remaining variant is above a threshold. We used 1 × 10^−4^ as a threshold and found no differences in model selection for a more lenient and more computationally expensive threshold of 1 × 10^−3^.

### 2.5. Test statistic and significance thresholds

Starting from the full final model, 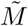, a test statistic for each variant *v* in the model is obtained by removing that single variant from 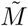 to obtain a new model 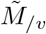. All of the variant’s associations are removed; thus, if 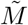 has *|M |*non-zero ***β*** parameters and a variant *v* in the model has associations with *P*_*v*_ phenotypes, the new model 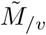 has *|M | – P*_*v*_ non-zero ***β*** parameters. The score *S*_*v*_ for a particular variant *v* is calculated as expected,

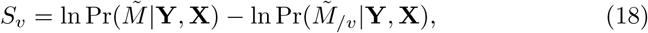

and serves as the test statistic for that variant. As mentioned above, the model 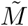 is optimal with respect to forward selection but not necessarily to backward selection. While the search biases *S*_*v*_ to positive values, negative values are possible and are observed, albeit infrequently.

Genome-wide randomizations were used to calibrate the value of *S*_*v*_, controlling for either family-wise error rates (FWER), with FWER = 0.05 corresponding to genome-wide significance, or false discovery rate (FDR), controlling for FDR = 0.05 as the threshold. Thresholds for FDR require fewer permutations for estimation; FDR was therefore used for simulated data and CHARGE data, whereas FWER was used for analysis of ARIC data. Permutations of the original data were generated starting with the vector of genotypes and the vector of phenotypes for each individual, with covariates already regressed out. Permutations were performed by randomizing the pairing between a genotype vector and its phenotype vector. Elements within individual genotype and phenotype vectors were not permuted. These permutations maintain the genetic covariance and phenotype covariance structure of the data.

Filtering steps independent of genotype-phenotype pairing, primarily filtering based on allele frequency and Hardy-Weinberg equilibrium, were identical for all permutations. Subsequent processing of permuted data sets exactly matched processing of the original data, including the computationally expensive step of performing all the genome-wide univariate tests. For FWER, the null distribution of the test statistic was obtained from 100 genome-wide permutations. The score of the best variant was retained for the 100 permutations, and the 5^th^-best score was used to define the 0.05 FWER threshold. For FDR, 10 permutations were done in the same way. The expected number of false discoveries 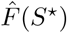 with scores greater than or equal to threshold *S*^*\**^ was estimated as

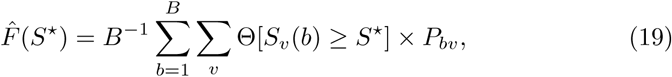

where *B* = 10 is the number of permutations, *S*_*v*_(*b*) is the score of variant *v* in permutation *b*, Θ(*u*) = 1 if logical argument *u* is true and 0 if false, and *P*_*bv*_ is the number of phenotypes associated with variant *v* in permutation *b*. The FDR for the unpermuted data for a given threshold was then calculated as

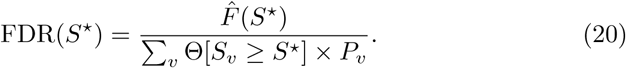

Variants from the unpermuted data were arranged in decreasing order by score; the first SNP *v*′ for which FDR(*S*_*v′*_) > 0.05 was identified; and the previous SNPs with *S*_*v*_ *> S*_*v′*_ were retained as the predicted positives at FDR 0.05. As is standard for methods based on model scores, calibration by permutation test is done separately for each data set analyzed.

### 2.6. Other methods

Assessments involved representative available implementations of major classes of pleiotropy methods. In general, adjustable parameters were set using published recommendations. We restricted attention to methods with run times that were small multiples of the cost of performing all *T* × *P* univariate tests of individual variants and phenotypes.

For aggregating association signals over a set of phenotypes, the methods SHom and SHet, using homogeneous and heterogeneous test statistics, were selected [2]. Given summary statistics of all *T* × *P* univariate tests (regression coefficients and standard deviations), SHom uses a generalized inverse-variance weighted analysis to combine individual tests into a pooled *z*-score, which also considers the correlation between phenotypes and sample sizes of different studies. By comparing the test statistic with a standard normal distribution, SHom obtains one p-value for each tested variant. SHet has a scoring function similar to SHom, but also introduces a threshold *τ*. The p-value for the test statistic combining all *t*-scores greater than *τ* is maximized, and the corresponding pvalue is assigned to the given SNP. Because of this selection process, SHet does not follow a standard normal distribution, and p-values are obtained through permutation.

For tests involving linear combinations of phenotypes, the Principal Components of Heritability (PCH) method [5] and MultiPhen [6] were selected. With PCH, a phenotype loading vector *w* is first estimated for each SNP. The loading vector is selected to maximize the variance of the loaded phenotypes explained by the given SNP using a subset of the data. Next, a *t*-score is obtained by regressing the genotype data onto the loaded phenotypes using the remainder of the data. By adopting a bagging technique and running cross-validation, the *t*-score distribution is estimated and a p-value is obtained.

For MultiPhen, each genotype is treated as a response variable outcome, and phenotypes are predictors. MultiPhen uses proportional odds logistic regression to regress genotype on the hyperplane constructed by the phenotypes, which models genotype data as ordinal [6]. No distributional assumptions are required for phenotypes, allowing MultiPhen to accommodate both binary and continuous measurements in a single framework. While ordinal regression for integer-valued allele dosages is theoretically attractive, the computational cost is much greater than for linear regression (Table 1). Furthermore, allele dosages estimated from imputation are real-valued rather than integer-valued. MultiPhen includes a gaussian kernel for this reason, and gaussian regression was used instead of ordinal regression for some of the results reported here. MultiPhen has two additional modes, variable selection and variable non-selection. For variable non-selection, all phenotypes are used as predictors for the genotype data; for variable selection mode, backward selection were performed on the phenotypes in order to exclude the phenotypes that were not associated with the SNP, and then the selected phenotypes were regressed on the genotype data. Non-selected phenotypes have nominal p-values of 1 as output.

**Table 1:** Running time for different methods for a simulation with 10,000 individuals, 6 phenotypes, and 638 SNPs of which 24 had associations, with Core i5 2.9 GHz CPU, 8 GB RAM. 12 SNPs are associated the first three phenotypes, while the other 12 SNPs are associated with the second three phenotypes.

| Method | Run Time |
| --- | --- |
| SHOM | 0m2s |
| SHET | 0m13s |
| SAPPHO-I | 1m1s |
| MultiPhen NonSelection Gaussian | 2m2s |
| PCH | 3m38s |
| SAPPHO-C | 4m39s |
| MultiPhen Selection Gaussian | 4m55s |
| MultiPhen NonSelection Ordinal | 30m15s |
| MultiPhen Selection Ordinal | 115m28s |

**Table 2:** Assessed methods provide qualitatively different types of predictions. ‘Parsimonious’ indicates that a single SNP is selected from an LD block, whereas ‘Non-Parsimonious’ indicates that all SNPs in an LD block are reported. ‘Phenotype selection’ indicates that the subset of associated phenotypes is reported, whereas ‘No phenotype selection’ indicates that the results do not specify which phenotypes are associated with a SNP and which are not.

|  | Parsimonious | Non-Parsimonious |
| --- | --- | --- |
| Phenotype selection | SAPPHO-I<br>SAPPHO-C | Univariate<br>MultiPhen-Selection |
| No phenotype selection |  | SHOM, SHET, PCH<br>MultiPhen-NonSelection |

We attempted to assess a regularized regression method, using an available implementation of the graphical fusion LASSO [8]. The shrinkage behavior of graphical fusion LASSO method requires optimization of regularization parameters *λ* and *γ* by cross-validation over a search grid. The cross-validation steps were computationally intensive, and for many (*λ, γ*) grid values the iterations did not converge. No other implementations for GWAS were readily available. Published results based on graphical LASSO are generally for smaller data sets; the available implementation was originally developed for a data set of 34 SNPs and 543 individuals [8]. We therefore excluded LASSO-based methods from the comparison.

Calibration of methods for 0.05 FWER or 0.05 FDR were conducted as for SAPPHO using the same set of genome-wide permutations. This calibration was conducted even for methods providing a nominal p-value based on assumed parametric distributions to ensure accurate benchmarking.

### 2.7 Assessment with real data

We assessed SAPPHO using individual-level phenotype and genotype data from the the Atherosclerotic Risk In Communities (ARIC) study cohort [20], focusing on phenotypes related to the electrocardiogram (ECG) parameters PR, QRS, and QT, which are risk factors for cardiovascular disease, sudden cardiac death, and stroke. This cohort includes approximately 8000 Caucasian ethnicity subjects and 2000 African-American ethnicity subjects. Assessments here use only the Caucasian ethnicity because power has been insufficient for AfricanAmerican ethnicity.

Known positive phenotype-genotype associations were taken from metaanalyses conducted by the CHARGE consortium, which includes ARIC as a cohort. Recent meta-analyses have included 88,000 individuals for PR, 40,407 for QRS, and approximately 100,000 for QT [13, 14, 15, 16]. Covariates for the EKG phenotypes were selected to be identical to those used in meta-analysis: height, age, gender, center, BMI, centerm, and heart rate.

Genotypes for ARIC were imputed by pre-phasing with Shapelt (v1.r532) and then imputing to 1000 Genomes [21] using IMPUTE2 [22, 23]. Measured SNPs used for imputation were restricted to MAF > 0.005, > 95% complete, HWE > 0.00001, resulting in 711,589 SNPs in the final set used for the imputation. Final imputations from IMPUTE2 used the reference panel 1000 Genomes haplotypes – Phase I intergrated variant set release (v3) in NCBI build 37 (hg19) in chunks of size 5Mb. All 1092 individuals were used for imputation from the reference panel.

Analyses were done focusing on the overlapping variants between ARIC and CHARGE cohorts, and we have tested that this overlap set of variants essentially includes all SNPs reported as significant by previous GWAS for ECG traits. Variants were removed if the ARIC or CHARGE genotypes violated Hardy-Weinberg equilibrium (*p* < 0.00001), were poorly imputed (Qual < 0.3), or if the minor alleles were too low frequency to have power (MAF < 0.01), corresponding to fewer than 160 copies of the minor allele. These criteria and filtering variants to require univariate p-value ≤ 10^−4^ and low LD with more significant variants (see Methods) resulted in 620 total variants for the PR, QRS, and QT phenotypes. This filtered list was used for each method.

Assessments were performed by defining variant-phenotype associations present in the meta-analyses at genome-wide significance (p-value ≤ 5 × 10^−8^ for each phenotype) as known positives. Assessments are complicated by linkage disequilibrium within the genome, which can lead to genome-wide significant findings for multiple variants within a linkage disequilibrium block. These multiple variants often correspond to a single causal variant, and for purposes of assessment they were grouped into a single known positive.

We performed the grouping as follows. For each phenotype, we linked together pairs of genome-wide significant SNPs within 500 kbp of each other. We then identified the distance-based connected components defined by these pairwise links. Each connected component in principle could contain multiple independent causal effects. To determine whether independent effects were present, we provided the SNPs in each connected component as a single locus to GWiS [10]. GWiS was then run separately on each connected component to select candidate SNPs representing independent effects.

In regions with strong association signals, these candidate sets may still contain more SNPs than independent effects. Furthermore, independent effects must be matched across phenotypes. We therefore used linkage disequilibrium as defined by *r*^2^ correlation to identify a final set of independent effects. We introduced correlation-based links between pairs of SNPs with *r*^2^ ≥ 0.05 and identified the connected components defined by the correlation-based links. This resulted with 107 gold-standard connected components, each connected component corresponding to a single effective known positive with one or more phenotypes.

We also investigated the robustness of the gold-standard connected components with respect to the *r*^2^ threshold of 0.05. For a threshold of *r*^2^ ≥ 0.01 the number of connected components was 90, and for *r*^2^ > 0.1 the number of connected components was 112. At the lower threshold, multiple effects are merged into a single connected component, while at higher threshold, one single effect may be divided into multiple connected components. With *r*^2^ ≥ 0.05, the SCN5A-SCN10A locus was assigned 3 different association signals, while at *r*^2^ ≥ 0.1 the SCN5A-SCN10A locus had 5 independent effects, which bracket the estimates from existing literature [13, 14, 15, 16]. While the details of performance of individual methods depend somewhat on the *r*^2^ threshold, the relative performance of different methods is stable with reasonable choices for the clustering threshold. Therefore for the current study all results were reported based on *r*^2^ ≥ 0.05 (Supplementary table 1).

Methods differ in their treatment of LD blocks and their attempt to identify the subset of phenotypes associated with each variant. The SAPPHO-I and SAPPHO-C methods attempt to provide a parsimonious list of associations, with only one SNP in each significant LD block. Other methods report each SNP within an LD block as a positive. The SAPPHO-I, SAPPHO-C, MultiPhen-Selection, and univariate methods identify the subset of phenotypes associated with a SNP, whereas other methods do not. For purposes of assessment, we calculated the *r*^2^ for each SNP selected by a method to each SNP in the gold standard correlation-based connected components. We defined *r*^2^ ≥ 0.1 as the threshold for matching. Each correlation-based connected component with at least 1 matching SNP was counted as a true positive; the remaining connected components were counted as false negatives. We tried different threshold including *r*^2^ ≥ 0.1, *r*^2^ ≥ 0.5, *r*^2^ ≥ 0.7, and *r*^2^ ≥ 0.8, and found that while using the thresholds other than *r*^2^ ≥ 0.1 did not substantially increase the number of true positives, they yielded many false positives, primarily non-causal variants somewhat correlated with real effects. We therefore used *r*^2^ ≥ 0.1 as the threshold for reporting results.

To be more favorable to the non-parsimonious methods, we used a similar grouping strategy to define the number of false positives. SNPs reported by a method but with *r*^2^ < 0.1 to any SNP in a gold-standard connected component were grouped into false-positive connected components using *r*^2^ ≥ 0.05, and each false-positive connected component was then counted as a single false positive. For SAPPHO, MultiPhen-Selection and univariate tests, the methods which provide the subset of phenotypes associated with each variant, we performed subsequent analyses to assess the ability to detect the correct variant-phenotype associations.

In addition to performing assessments with the original ARIC data, we also performed assessments in which the ARIC data was augmented with with random phenotypes generated as independent and identically distributed standard normal random variables. We performed tests with 3, 6, and 10 random phenotypes added to the 3 ECG phenotypes. These assessments were designed to identify robustness of methods when phenotypes with shared genetic factors are unknown.

SAPPHO was then run in summary mode using sufficient statistics from CHARGE analyses. The sufficient statistics included phenotype covariances, phenotype-genotype regression coefficients and standard errors, allele frequencies, genotype-genotype covariances, and sample numbers for each phenotype-genotype association. For SAPPHO-C, the difference in sample numbers complicated the likelihood ratio term in the score statistics, and the computational expense increased dramatically as more associations were included into the model; therefore, only SAPPHO-I was run in this case. Permutations were performed by randomly resampling 100,000 individuals from the ARIC primary data. To be more specific, individuals from ARIC data were resampled with replacement for 100,000 times to construct 100,000 ‘new’ individuals. With this procedure, the genotype allele distribution for resampled population should be consistent with that of the ARIC population. Shuffling and subsequent steps were then performed exactly as for the ARIC primary data to preserve the underlying phenotype-phenotype correlation was preserved. Results for 0.05 FDR used 10 population-wide, genome-wide permutations. The methods for constructing the gold standard and assessing true and false positives were the same as for the ARIC primary data.

To assess the biological relevance of new associations identified as significant by SAPPHO for the CHARGE cohort, enrichment analysis was performed for genes detected by SAPPHO-I at 0.05 FDR. The analysis focused on the curated gene sets including BIOCARTA, KEGG, REACTOME, and GO pathways as aggregated by MSigDB [24]. For this analysis, a 2 × 2 contingency table was constructed for each pathway, with 0/1 columns denoting whether the gene was detected by SAPPHO-I at 0.05 FDR as the columns, and 0/1 rows denoting whether that gene was in the pathway. Fisher’s exact test was then run on each contingency table to obtain a one-sided p-value for a one-sided test of enrichment. We performed this assessment first for all the loci reported by SAPPHO-I. We then modified the procedure to account for the possibility that some of the pathway assignments found in MSigDB may have been influenced by the GWAS contributing to CHARGE, whose data we are using. Our modification was to exclude all gold-standard loci from consideration, removing them both from the SAPPHO-I results and from the gene sets. We then performed 2×2 contingency enrichment analysis as before but restricted to the non-gold-standard loci.

### 2.8. Assessment with simulated data

Methods were run on simulated data to gain further insight into differences in performance due to controlled aspects of genetic and environmental architecture of complex phenotypes. Simulations followed previous protocols used to assess GWiS and other gene-based tests [10]. Simulated data sets included 1,000,000 independent SNPs for 10,000 individuals. Minor allele frequencies for each SNP were generated uniformly between 0.01 and 0.5 to model common variants. Three sets of simulations were done using frameworks denoted ‘genes only’, ‘genes and environment’, and ‘genes only with random phenotypes’. Each phenotype *y* with genotype vector **x** was simulated as

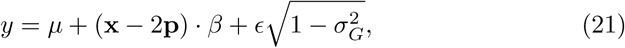

where *µ* is the overall phenotype mean, **p** is the vector of minor allele frequencies, *β* is the vector of regression coefficients for the scenario, *ϵ* is a unit normal random variable, and 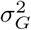 is the genetic variance,

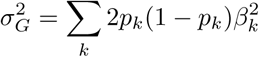

where *p*_*k*_ denotes the minor allele frequency of SNP *k*. The entry *β*_*k*_ = 0 if SNP *k* is not associated with the phenotype. For an associated SNP,

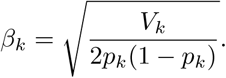

The term *V*_*k*_ is chosen based population size *N* and specified type I and type II error rates *α*_1_ and *α*_2_ for univariate tests as

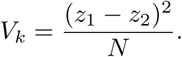

For a two-tailed test, *z*_1_ = Φ^−1^(1 – *α*_1′_2). The term *z*_2_ is set by the desired type II error rate *α*_2_ *≡* 1 – power as *z*_2_ = Φ^−1^(*α*_2_). The function Φ^−1^ is the inverse of the standard normal cumulative distribution with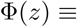 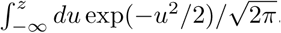. With this setting, a SNP with regression coefficient *β*_*k*_ will have the specified power at threshold *α*. All simulations were performed 5 independent times, including 10 sets of permutations each to determine the 0.05 FDR threshold.

In the ‘genes-only’ scenarios, phenotypic correlations were due entirely to shared genetic variants with no environmental effects. These simulations considered 6 phenotypes and 24 SNPs with causal effects. Simulations were performed separately for 3 scenarios reflecting increased sharing of genetic factors: independent, in which 4 SNPs were associated with each phenotype and no SNPs shared between phenotypes (24 total pairwise SNP-phenotype associations); block, in which the 6 phenotypes were divided into 2 blocks, and each block was associated with 12 SNPs (72 total SNP-phenotype associations); and full, in which the 24 SNPs contributed to each of the 6 phenotypes (144 total SNP-phenotype associations). Effects for all SNPs were set to have 50% power at 5 × 10^−8^ threshold.

For ‘genes-and-environment’ simulations, phenotypic correlations were due to both genetic and environmental effects. Two scenarios were simulated in this case: weak environment and strong environment. For weak environment, 4 phenotypes were partitioned into 2 blocks of 2 phenotypes; phenotypes within the same block are correlated through environmental effects, while phenotypes across different blocks are not; for strong effects, the 4 phenotypes are all correlated through environmental effects. For these scenarios, the random variable ϵ for each phenotype *p* (Eq.21) follows a multivariate normal with Var(ϵ_*p*_) = 1 and with covariance Cov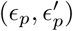 determined by a predefined environmental co-variance structure. For the weak environment simulation, the 4 × 4 covariance matrix is a block matrix of two matrices of size 2, and for strong environmental simulation, all elements of the 4×4 matrix are non-zero. For this study, we chose to set the correlation for different environmentally correlated phenotypes to 0.5. For weak environment, SNPs were associated with the phenotypes in 4 modes: different-block-same-effect, where each SNP is associated with two phenotypes, one from each block, and the effects are in the same direction; different-blockdifferent-effect, where each SNP is associated with two phenotypes, one from each block, and the effects are in opposite directions; same-block-same-effect, where each SNP is associated with two phenotypes from the same block, and the effects are in the same direction; same-block-different-effect, where each SNP is associated with two phenotypes from the same block, and the effects are in opposite directions. For strong environment, SNPs were associated with the phenotypes in 2 modes: same effect, where SNPs are correlated with all phenotypes with effects of the same direction; different effect, where SNPs are correlated with first two phenotypes with positive effects, and the last two phenotypes with negative effects. Different directions of effect were represented by different signs for the regression coefficients, with magnitudes defined based on type I and type II error rates exactly as in the ‘genes-only’ simulation. All effects were simulated to have 50% power at 5 × 10^−8^ univariate test threshold. For weak environment, 24 SNPs were simulated and divided equally into 4 modes, with 6 SNPs in each mode, ending up with 48 total associations; for strong environment, 24 SNPs were simulated and divided equally into two modes, with 12 SNPs in each mode, ending with 96 total associations. With this set of simulation assessed the capability of different methods to detect SNPs whose association patterns have the same or opposite direction from the environmental association patterns.

For the ‘mixture of genetic and non-genetic phenotypes’ simulations, we attempted to generate scenarios similar to the real ARIC data with known association patterns. Therefore, 13 total phenotypes were simulated, and the simulations were done with three scenarios: ONE association, where all active SNPs were associated with phenotype 1; TWO associations, where all active SNPs were associated with phenotypes 1 and 2; and THREE associations, where all active SNPs were associated with phenotypes 1, 2, and 3. For each scenario, all active SNPs follow the same association pattern, with the number of associated phenotypes differing between the scenarios. The effect of each association was simulated such that each SNP has 50% power at 5 × 10^−8^ threshold, with the two associations and three associations effect calculated as follows using *P* to denote power:

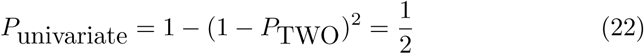

which gives us *P*_TWO_ = 0.293.

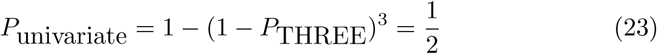

which gives us *P*_THREE_ = 0.206. 24 SNPs were simulated to be associating variants in each scenario, yielding 24, 48, and 72 total associations.

## 3 Results

### 3.1. Comparison of performance for different methods on ARIC data

For evaluation with primary data, both SAPPHO-I and SAPPHO-C, together with other pleiotropy methods, were applied to identify shared genetic contributions to ECG traits (Fig. 2). Methods were calibrated using permutations to identify the appropriate threshold corresponding to a 0.05 FWER for genome-wide tests of 3 phenotypes. The conventional threshold for univariate tests of 3 independent phenotypes would be (5*/*3) × 10^−8^ = 1.67 × 10^−8^. Calibration by permutation for univariate tests gave a single-test threshold of 1.068 × 10^−8^. With ARIC data, SAPPHO-C was able to recall 15 known loci, better than any other methods, followed by SAPPHO-I, finding 13 known loci (Fig. 2). The MultiPhen and SHet methods are next best in performance, returning 8 or 9 true positives, but no better than standard univariate tests. The methods PCH and SHom perform worse than univariate tests (Fig. 2, Supplementary Table 2).

**Figure 2:**
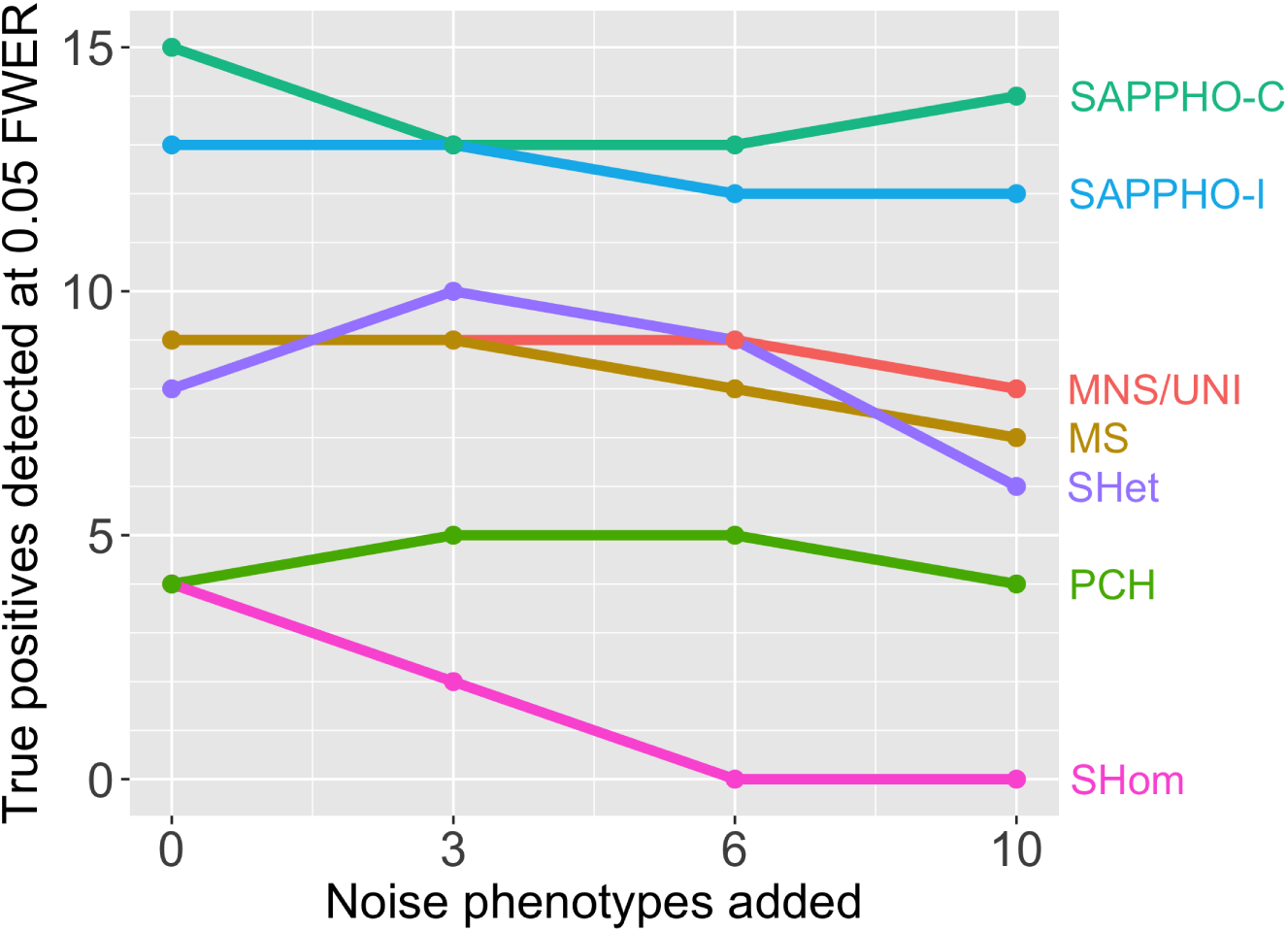
Pleiotropy methods were used to detect associations with the PR, QRS, and QT phenotypes in the ARIC cohort. The three measured phenotypes were then augmented with 3, 6, and 10 noise phenotypes. SAPPHO had the greatest power to detect associations regardless of noise phenotypes were present. The pleiotropy methods SHet, MS, MNS had power similar to standard univariate tests. PCH and SHom had lower power than univariate tests, and the performance of SHom degraded further as noise phenotypes were added. All methods were controlled for type I error at FWER = 0.05. Methods were MultiPhen-Selection (MS), Mutiphen-NonSelection (MNS), Principal Components of Heritability (PCH), Homogeneous Test Statistics (SHom), Heterogeneous Test Statistics (SHet), and univariate tests corrected using permutations (UNI).

We then investigated the variants identified by each method. The set identified by SAPPHO included the variants identified by all other methods, except for a single locus found by SHet. This SNP, rs1896312, has known positive associations with *P*_*CHARGE_PR*_ = 1.151 × 10^−34^ and *P*_*CHARGE_QRS*_ = 2.626 × 10^−9^.P-values from ARIC for univariate tests were *P*_*ARIC_PR*_ = 1.5 × 10^−6^ and *P*_*ARIC_QRS*_ = 3.3 × 10^−3^. The p-value returned by SHet was 2.94 × 10^−9^, better than its FWER=0.05 threshold value 1.52 × 10^−8^. The SNP was added to both SAPPHO-I and SAPPHO-C models, but did not pass the 0.05 FWER threshold. The reason that this loci was only detected by SHet is that as observed with ARIC data and simulated data, SHet is powered to find variants that have weak associations across all or most of the given phenotypes. For SAPPHO, given its stringency for adding associations to the model, only the PR association was detected, and thus, this SNP was not reported as being significant.

Two SNPs were identified by multiple methods, yet were not in the goldstandard: rs7638275 and rs17608766. rs7638275 has *P*_*CHARGE_PR*_ = 0.845, *P*_*CHARGE_QRS*_ = 0.41, and *P*_*CHARGE_QT*_ = 0.4667, and was therefore not included in the gold-standard. However, it has *P*_*ARIC_PR*_ = 5.27 × 10^−13^ and *P*_*ARIC_QRS*_ = 1.97 × 10^−6^, strong evidence for a true association within the ARIC subpopulation. We note that rs7638275 was reported as a rare variant with low imputation quality for most cohorts in CHARGE, and therefore it was not detected for any of the three ECG traits. In ARIC, however, rs7638275 was well imputed with a 1.5% minor allele frequency, and therefore detected by methods including SAPPHO-I, SHet, MultiPhen, PCH, and univariate tests. SAPPHO-C did not identify rs7638275 because its effect was partially explained by correlated SNPs already in the model. We reached this conclusion by attempting to add rs7638275 to the SAPPHO-C model; we found that its p-values were much less significant, 2.11 × 10^−5^ for PR and 0.01745 for QRS.

The other SNP, rs17608766, had p-values *P*_*CHARGE_ PR*_ = 1.7 × 10^−7^ and *P*_*CHARGE_ QRS*_ = 1.2 × 10^−5^ for CHARGE, and *P*_*ARIC_ PR*_ = 1.4 × 10^−7^ and *P*_*ARIC*_*_* _*QRS*_ = 3.0 × 10^−4^ in ARIC cohort. It was not included the gold standard because none of its associations passed the p-value 5 × 10^−8^ threshold. This SNP was found by both SHom/SHet and SAPPHO-I using data from ARIC cohort. This SNP was also later found by SAPPHO-I run on CHARGE metaanalysis results, which is strong evidence for its association with ECG traits (Supplementary Table 3). This finding was supported by a recent study which reported rs17608766, located in the gene *GOSR2*, to be associated with cardiac structure and function [25], which exhibits SAPPHO’s capability to identify novel pleiotropic associations.

### 3.2 Pleiotropy of ECG traits in ARIC and CHARGE

SAPPHO-I was run in summary mode on CHARGE meta-analysis summary results to see whether any additional pleiotropic SNP could be identified. The other methods were not run with CHARGE summary results, for two different reasons: PCH and MultiPhen require primary phenotype and genotype data which are not available for CHARGE, and SHom/SHet perform poorly based on results from ARIC cohort and simulations. These results, together with the association pattern obtained from ARIC and the gold-standard, are shown in (Fig. 3).

**Figure 3:**
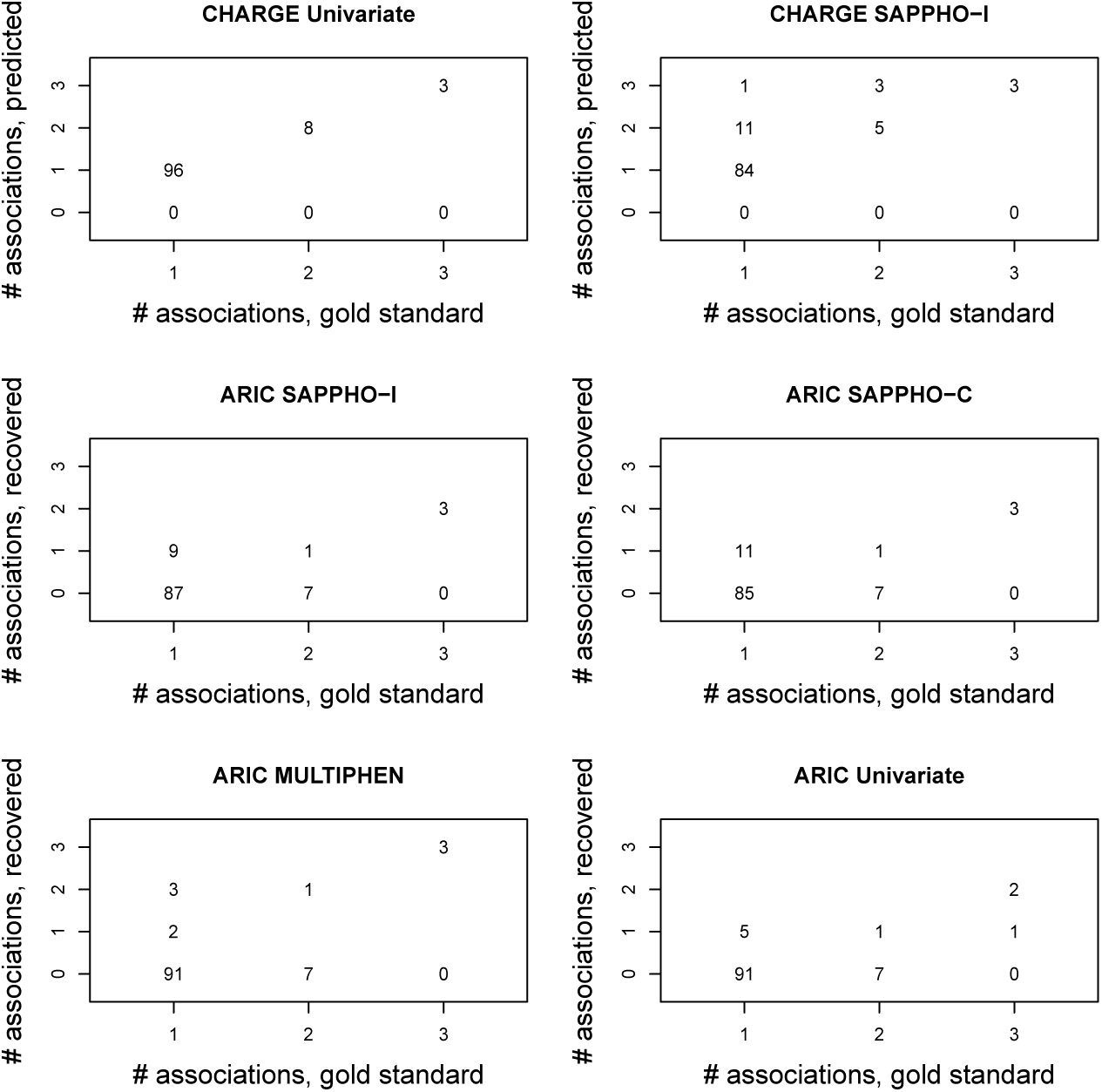
Number of associations recovered versus number previously known (*p* < 5 × 10^−8^ in CHARGE for univariate single phenotype test). First row: CHARGE meta-analysis data as analyzed by univariate tests for each phenotype (*p* < 5 × 10^−8^) and by SAPPHO-I (0.05 FDR). The first row denotes univariate test (with *p* < 5 × 10^−8^ cut-off) and SAPPHO-I run on CHARGE meta-analysis results; Second row: ARIC data as analyzed by SAPPHO-I and SAPPHO-C using 0.05 FWER threshold. Third row: ARIC data as analyzed by MultiPhen and univariates test using 0.05 FWER threshold, equivalent to *p* < 5 × 10^−8^*/*3 for univariate tests. More discoveries are made with CHARGE (top row) because it is a larger cohort and because 0.05 FDR is a less stringent threshold than 0.05 FWER. SAPPHO has greater power for the ARIC cohort than MultiPhen or univariate tests. Pleiotropic associations discovered by MultiPhen in the ARIC cohort (bottom left panel) may be over-predictions as these associations were not genome-wide significant in the much larger CHARGE cohort.

The genetic architecture of ECG traits includes SNPs contributing to distinct subsets of PR, QRS, and QT phenotypes. Given that ARIC is a subset of the CHARGE cohort, the power to detect true associations using ARIC data is smaller compared to using the entire CHARGE data. Therefore, for SAPPHO and univariate tests run on ARIC, in some cases gold-standard SNPs were not detected at all; in other cases, the strongest associations of a SNPs are retained but other gold-standard associations are lost. As is seen in (Fig. 3), the predicted number of associations is smaller than the expected number of associations, and the numbers denoting count of real hits lie below the diagonal line. For MultiPhen run on ARIC, however, the number denoting real hit counts all lie on or above the diagonal line, indicating that more associations were found for certain loci in the gold standard. Given the much smaller power of ARIC compared to CHARGE, these are likely to be over-predicted associations from MultiPhen rather than true pleiotropic loci; this over-predicting behavior of MultiPhen was later observed in the simulation studies as well (Supplementary Table 4, association pattern).

With ARIC data, SAPPHO methods were able to retrieve more hits than either MultiPhen or univariate tests. For CHARGE data, SAPPHO-I run in summary mode was able to retrieve all the real hits in the gold standard. Additional associations were found with SAPPHO-I for some gold standard loci, yielding additional loci with pleiotropic effects (Table 3). The number of additional associations added depends on the tuning parameter; for this test we set the *γ* parameter to allow for associations with *p* < 10^−4^ to be added.

**Table 3:**
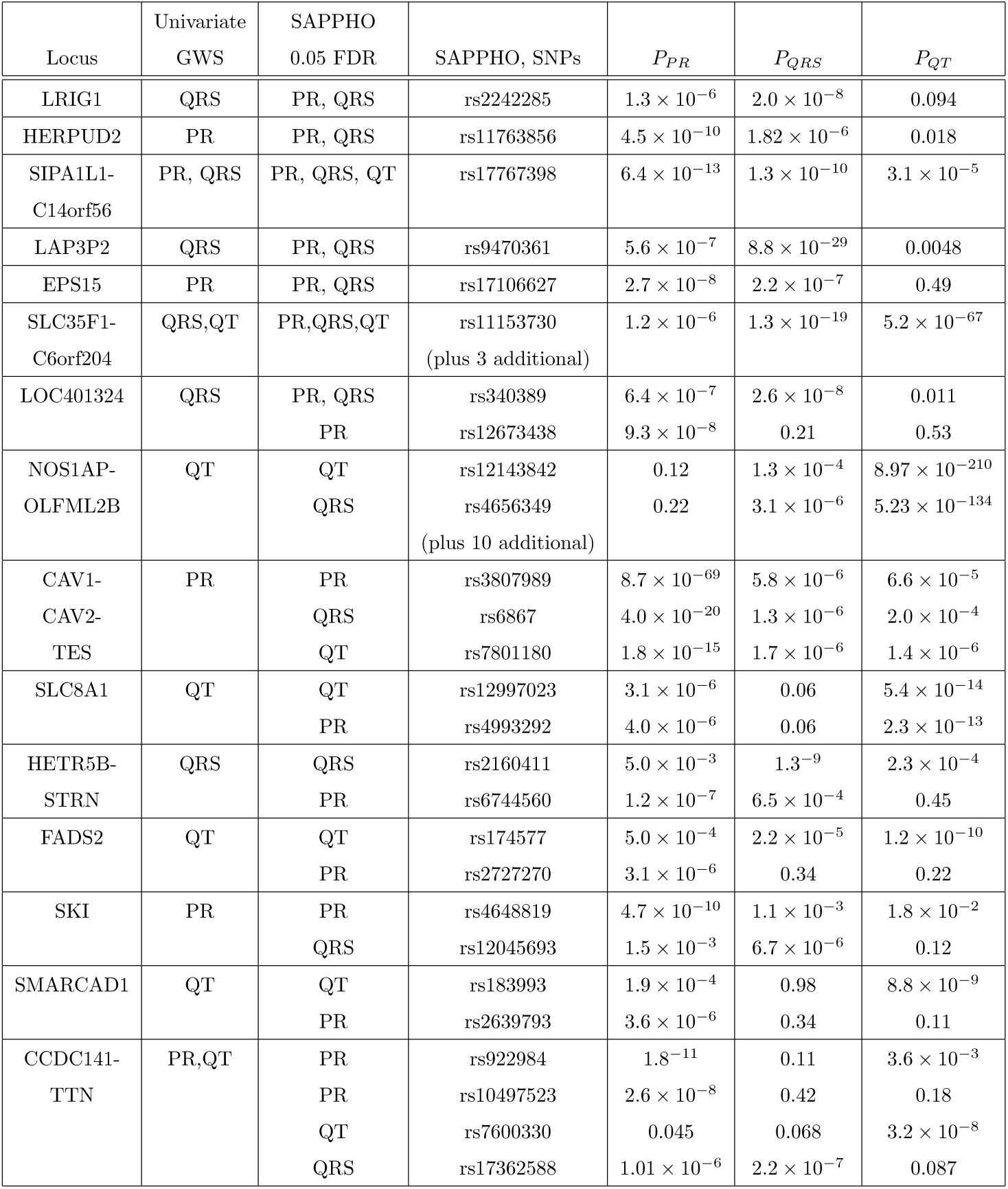
Additional novel associations detected by SAPPHO-I in known gold-standard loci. **Locus**: Gene symbol or symbols spanning an associated region. **Univariate GWS**: phenotypes detected at 5 × 10^−8^ genome-wide significance threshold. **SAPPHO 0.05 FDR**: phenotypes detected by SAPPHO at 0.05 FDR. **SAPPHO, SNPs**: SNPs detected by SAP-PHO, with rows corresponding to the previous *SAPPHO 0.05 FDR* column. ***P*_*P* *R*_, *P*_*QRS*_, *P*_*QT*_**: univariate p-values from CHARGE meta-analysis. New associations added to a locus are in two categories: (1) a variant already associated with at least one trait is associated with a new trait or traits (the first 7 loci in the table); (2) a new variant is introduced and is associated with a trait not previously associated with the locus (the last 8 loci in the table).

For loci already part of the gold standard, two types of new associations were added (Table 3): (1) a variant already associated with at least one trait was associated with at least one additional trait; (2) a variant not previously associated with any trait was associated with a new trait not yet associated with that locus. In loci not part of the gold standard, SAPPHO detected 124 new hits at 0.05 FDR. Most were associated with single traits, but the above-mentioned SNP rs17608766 in *GOSR2* was detected as pleiotropic. Pleiotropic effects at the locus level were observed more frequently. For example, *SLC12A7* contains rs2334955 and rs4285270, which were associated with QT and PR respectively, and *KLHL38* contains rs4871397 and rs16898685, which were associated with PR and QRS respectively. The linkage disequilibrium for each of these pairs of SNPs is weak, *R*^2^ = 0.064 for rs2334955 and rs4285270 and *R*^2^ = 0.0017 for rs4871397 and rs16898685, suggesting two independent effects within each locus. These associations reveal new genetic connections between different ECG traits.

Our analysis of CHARGE has no known ground truth; therefore, we used biological annotations to assess performance and gain insight. These assessments tested for enrichment of genes identified at 0.05 FDR for membership in annotated gene sets (see Methods). A total of 7246 gene sets were analyzed, corresponding to a nominal p-value of 6.9 × 10^−6^ for conventional significance. Genes detected by SAPPHO at 0.05 FDR show strong enrichment signals for several pathways involved with cardiac physiology and activities (Supplemental Table 5). The three most significant gene sets represent regulation of heart contraction (*p* = 2.5 × 10^−19^), muscle systems processes (*p* = 3.6 × 10^−19^), and cardiac conduction (*p* = 3.7 × 10^−19^). Additional notable categories include regulation of heart rate (*p* = 3.0 × 10^−17^), heart process (*p* = 1.1 × 10^−14^), and cardiac muscle cell action potential (*p* = 4.3 × 10^−14^). To ensure that these findings were robust, we repeated the analysis but excluded the significant findings from previously published GWAS. Gene sets specific to cardiac electrophysiology remain highly significant, including regulation of heart contraction (*p* = 3.5 × 10^−8^), muscle systems processes (*p* = 2.3 × 10^−9^), cardiac conduction (*p* = 1.6× 10^−7^), as well as heart rate (*p* = 5.9× 10^−6^), heart process (*p* = 5.9×10^−6^), and cardiac muscle cell action potential (*p* = 3.1×10^−6^). These findings support the conclusion that the novel loci with statistical associations detected by SAPPHO increase are indeed causal for ECG traits.

Given that pleiotropy is observed for ECG traits, we asked whether variants that affect the same subset of traits typically have the same direction of effect, and whether this depends on the observed phenotypic correlation. For ECG traits, Cor(PR, QRS) = 0.051, Cor(PR, QT) = -0.026, Cor(QRS, QT) = 0.168, where Cor(*x, y*) stands for the correlation between the two traits *x* and *y*. The consistency of direction of effects and phenotypic correlations for pleiotropic SNPs detected by SAPPHO on CHARGE are shown in Table 4, where ‘+/+’ denotes the SNPs whose effects are in the same direction, and ‘+/-’ denotes the count of SNPs whose effects are in different directions. Out of the 23 pairs of association effects, 16 were consistent with the phenotypic correlations. The probability that association effects were consistent with phenotype correlation was 0.70 with a 95% binomial distribution confidence interval of [0.49, 0.84]. We conclude that SAPPHO is able to detect variants whether or not the direction of genetic effect matches the overall direction of correlation. We further conclude that variants that contribute to the same pair of phenotypes can often show different directions of effect, suggesting that distinct biological mechanisms connect the variants to their downstream effects on ECG traits.

**Table 4:** Consistency of direction of effects and phenotypic correlations for each pair of ECG phenotypes. ‘Correlation’ denotes the correlation of each phenotype-phenotype pair; ‘+/+’ denotes the count of SNPs whose effects are in the same direction; ‘+/-’ denotes the count of SNPs whose effects are in different directions. Of the 23 pairs of association effects, 16 were consistent with the phenotypic correlation, corresponding to a probability of 0.70 (binomial parameter 95% confidence interval [0.49, 0.84]) that the direction of effect agreed with the phenotypic correlation. These results demonstrate that SAPPHO is able to detect variants whether or not the direction of genetic effect matches the overall direction of phenotypic correlation.

| Phenotypes | Correlation | +/+ | +/- |
| --- | --- | --- | --- |
| PR-QRS | 0.051 | 9 | 3 |
| QRS-QT | 0.168 | 3 | 4 |
| PR-QT | -0.026 | 0 | 4 |

### 3.3 Performance of different methods with additional null phenotypes

We also investigated performance of each method as noise was introduced through addition of null phenotypes (Fig. 2). This assessment models a collection of phenotypes where only small subsets share genetic factors, and these subsets are unknown at the outset. The least robust method is SHom, which makes the assumption that all traits share genetic factors. Other methods also lose power when noise phenotypes are presented, though to a lesser degree. For real-world application, variants are not likely to be associated with all inputed phenotypes, making robustness when noise phenotypes are presented crucial.

### 3.4 Genes-only simulations

Methods were assessed with phenotypes with shared genetic factors but without shared environmental contributions (Fig. 4). Three scenarios were considered, with increased sharing of genetic factors: independent, with 4 independent SNPs contributing to each of 6 phenotypes; block, with 12 SNPs contributing to one block of 3 phenotypes, and 12 other SNPs contributing to a second block of 3 phenotypes; and full, with 24 SNPs contributing to all 6 phenotypes. In general, all pleiotropy-based methods performed better with increasing shared genetic factors. Univariate tests were run and real-positives were determined with three methods: UNI where the 0.05 FDR threshold was used; UNILE with LE standing for loose-empirical where the standard 5×10^−8^ threshold were used for each association; UNISE with SE standing for stringent-empirical where the 5 × 10^−8^ threshold was corrected with the number of phenotypes tested, which in this case is 6.

**Figure 4:**
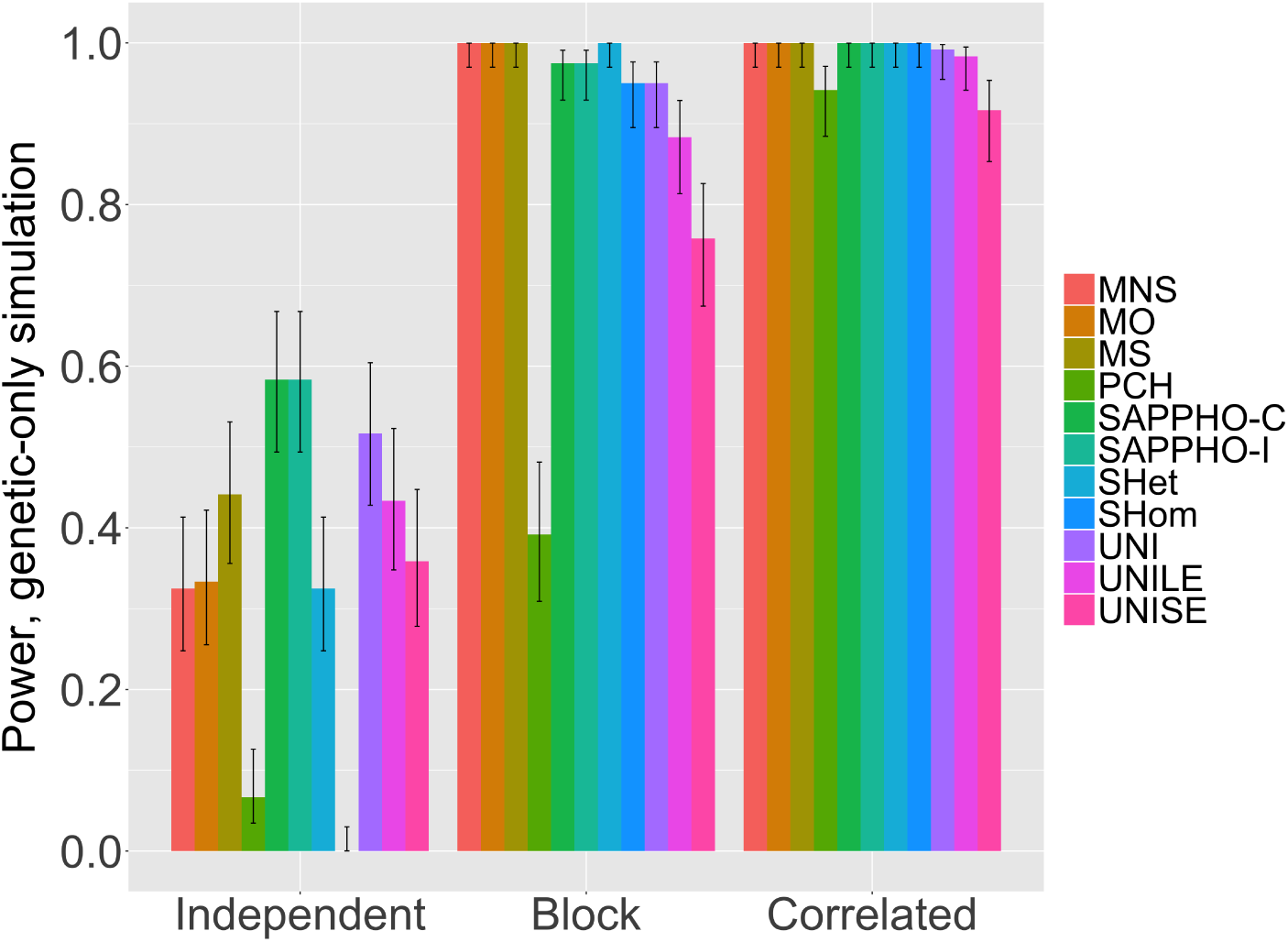
Power for associations of 6 phenotypes simulated to be correlated through genetic effects only. Scenarios are Independent (6 phenotypes, no pleiotropy), Block (2 blocks of 3 phenotypes each), and Correlated (all 6 phenotypes correlated). Error bars indicate 95% confidence intervals estimated from 5 repeated runs and a binomial distribution. For the independent scenario lacking pleiotropy, SAPPHO methods performed the best. For the block correlation scenario, MultiPhen leading performed best, followed by SAPPHO. For a single correlated block, all pleiotropy methods perform well. Methods were MultiPhen-NonSelection (MNS); MultiPhen Ordinal regression (MO); MultiPhen-Selection (MS); univariate test corrected with permutation (UNI); univariate test at loose empirical threshold *p* < 5 × 10^−8^ (UNILE); univariate test at stringent empirical threshold *p* < 5 × 10^−8^*/*6 for 6 phenotypes (UNISE).

**Figure 5:**
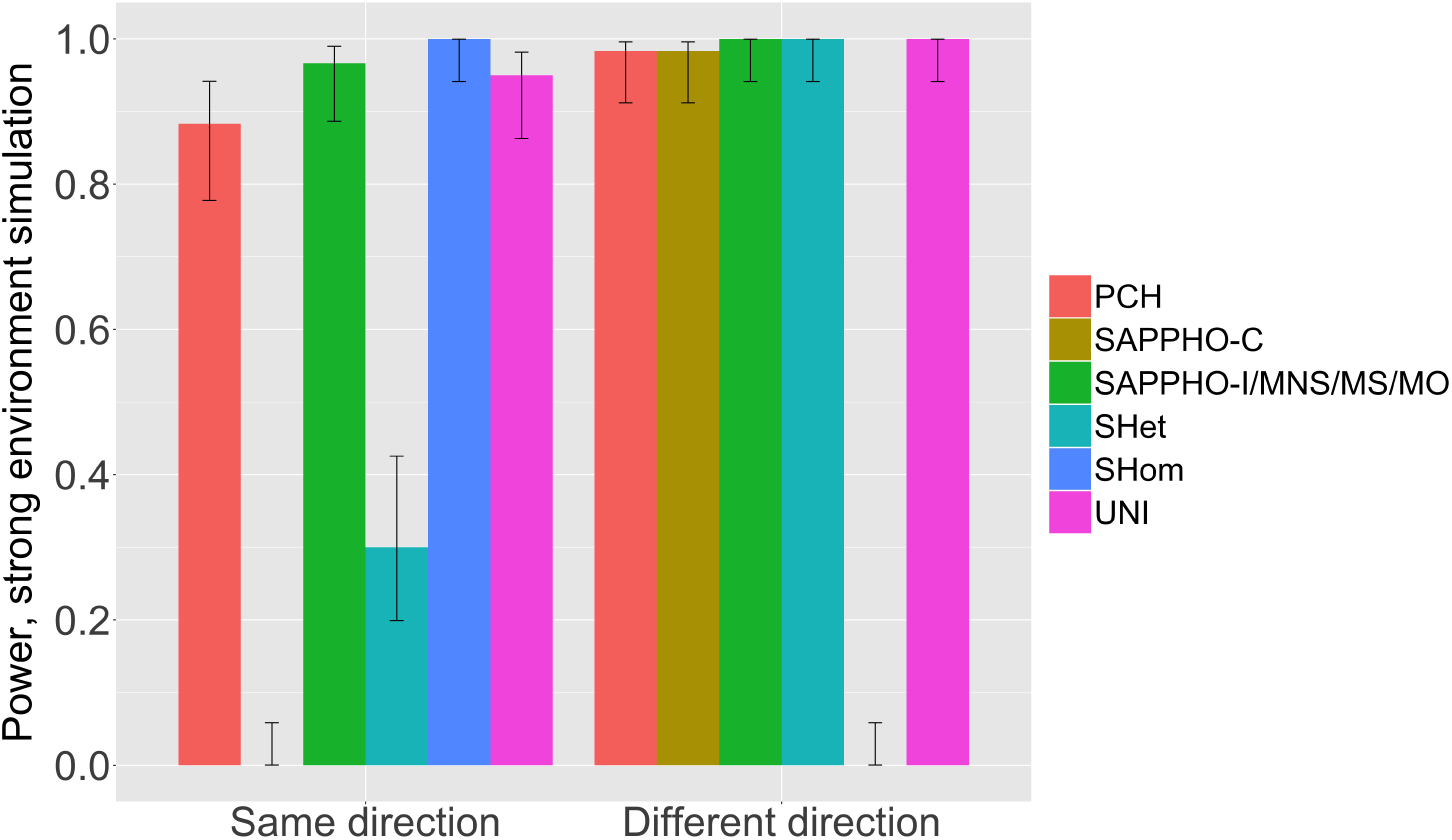
Power to detect associations for scenarios involving strong environmental correlations. Two sets of SNPs each containing 12 variants were simulated to contribute to 4 phenotypes. For the same-direction scenario, all 12 SNPs have positive effect with all phenotypes; for the different-direction scenario, the 12 variants had positive effects for the first 2 phenotype and negative effects for the second 2 phenotypes. SAPPHO-C and SHet were unable to detect same-effect SNPs; SHom could not detect different-direction SNPs. Methods were MultiPhen-NonSelection (MNS); MultiPhen Ordinal regression (MO); MultiPhen-Selection (MS); univariate test corrected with permutation (UNI).

**Figure 6:**
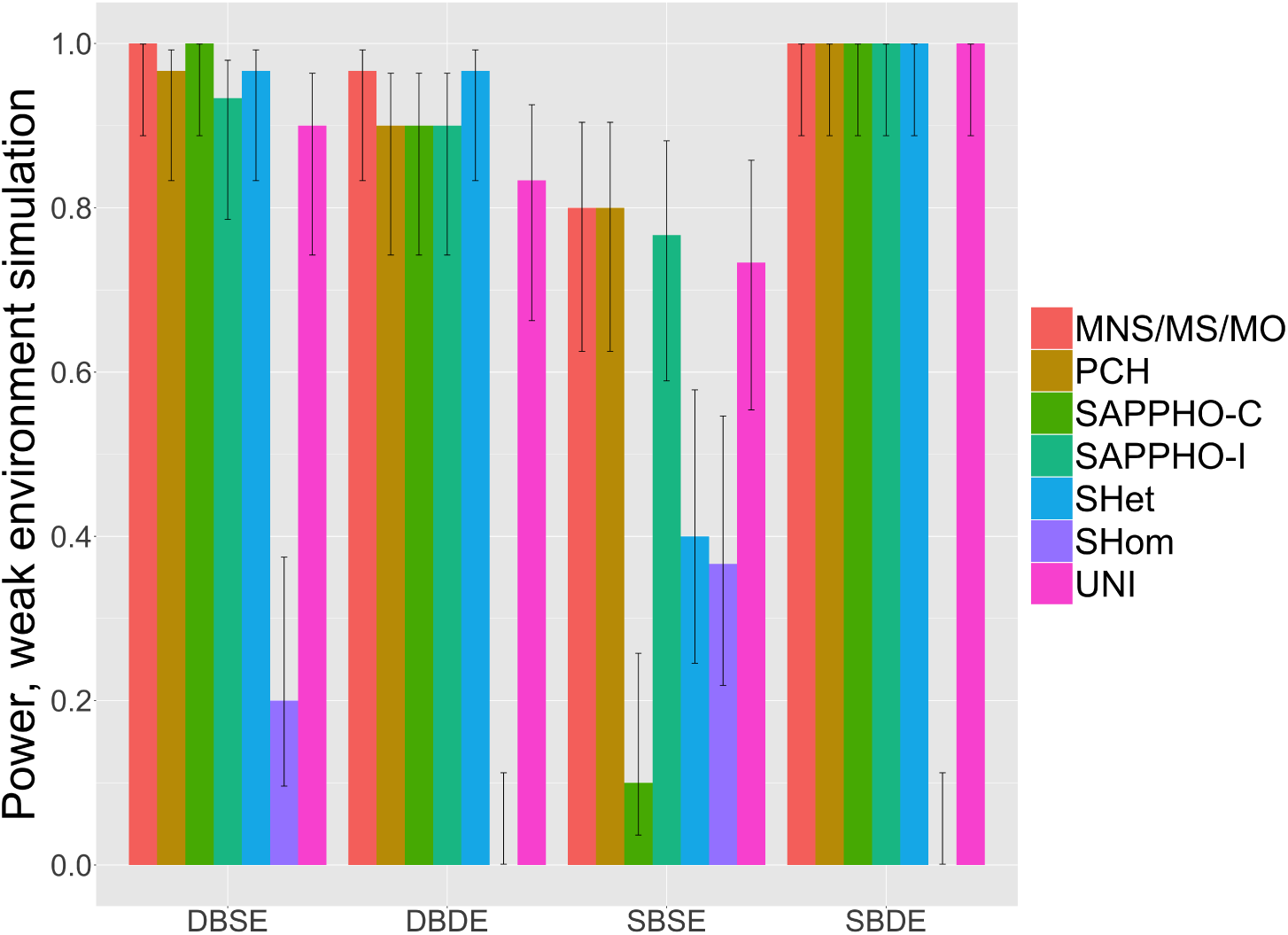
Power to detect associations for scenarios involving weak environmental correlations. Four sets of SNPs each containing 6 variants were simulated to contribute to 4 phenotypes. Scenarios as described in the main text were DBSE, different-block-same-effect; DBDE, different-block-different-effect; SBSE, same-block-same-effect; and SBDE, same-blockdifferent-effect. SAPPHO-I and MultiPhen performed the best over all scenarios. SAPPHOC experiences dramatic loss of power under the sbse scenario, similar to its loss of power for similar scenarios involving strong environmental correlations. Methods were MultiPhen-NonSelection (MNS); MultiPhen Ordinal regression (MO); MultiPhen-Selection (MS); univariate test corrected with permutation (UNI).

The performance of SAPPHO, assessed as power to detect at genome-wide significance, out-performed all other methods for the independent scenario. Methods other than SAPPHO have lower power than univariate tests when phenotypes do not share causal variants. For the full scenario, pleiotropy methods other than PCH and UNISE had power close to 100%, making the methods difficult to distinguish on this basis. For the block scenario, the MNS (MultiPhennon-select) and MS (MultiPhen-select) methods were somewhat better than other methods, outperforming SAPPHO-I, SAPPHO-C, and SHet by 1 to 2 hits in 2 out of 5 runs. These simulations suggest that SAPPHO’s superior performance on real data comes in part from robust performance even when pleiotropy is absent.

### 3.5 Genes-and-environment simulations

As described in the method section, for the genes-and-environment simulations, phenotypic correlations were set to be due to both genetic and environmental effects. Simulations were run in two modes: strong environment and weak environment.

For strong environment same-effect SNPs, most methods performed well except for SHet and SAPPHO-C, which were unable to detect most associations for this group of SNPs. SHom was able to detect all SNPs because the underlying association pattern was same as the simulated pattern, namely the variant is associated with all phenotypes. Although the underlying assumption for SAPPHO-I did not match the simulation pattern, it performed well because all the SNPs followed the same association pattern; thus the pattern prior gives its power to detect all the variants.

Unexpectedly, SAPPHO-C performed poorly in the same-effect setting. The reason is likely because the variance explained by one association is compensated through correlation with other phenotypes, with the result that adding a variant to the model does not improve the model score. To explore this effect further, we calculated the SAPPHO-C score for the true model and found it to have a large negative score. We can also explain this effect by noting that the BIC penalty assumes that regression coefficients for a SNP have independent sign, whereas the architectures in this scenario force the regression coefficients to have the same sign, resulting in a penalty that is too large.

For strong environment different-effect SNPs, all methods performed well, except for SHom, due to the difference in the simulated data and its underlying assumption.

For weak environment simulations, SNPs were simulated to follow 4 different patterns: different-block-same-effect(dbse), different-block-different-effect(dbde), same-block-same-effect(sbse), same-block-different-effect(sbde). Different/same-block denotes whether the environmental correlation is identical with the genetic pleiotropic effects, and same/different effect denotes whether the effects for the simulated pleiotropic SNPs are of the same direction or not. For dbse, SAPPHOC and MultiPhen performed the best by finding all associated SNPs; followed by SAPPHO-I, SHet, and PCH; for dbde, SAPPHO-C and MultiPhen again performed the best. This is somewhat similar to the genetic only simulation, where the correlation between phenotypes are only through genetic effect, and SNPs are correlated with both blocks. For sbse, SAPPHO-I performed the best, while SAPPHO-C, SHet, and SHom performed poorly; this situation is similar to the same-effect SNPs for strong environment simulation, where a SNP is associated with all phenotypes in a block, with positive effects. Similarly, the performance on sbde was same as different-effect for strong environment simulation, with all methods performing well. For both simulations, the original version of MultiPhen using ordinal regression was again performed, yielding outcomes similar to using Gaussian regression, but with much longer running time.

### 3.6. Mixture of genetic and non-genetic phenotypes

We performed simulations with genetic effects in 3 out of 13 total phenotypes to further investigate the relative performance of SAPPHO and MultiPhen and, in cases where SAPPHO performed less well, whether the cause was the underlying statistical model or the greedy rather than full model search. These simulations used three scenarios labeled ONE, TWO, and THREE. In scenario ONE, all associations were with phenotype 1. In scenario TWO, all associations were pleiotropic with phenotype 1 and phenotype 2. In scenario THREE, all associations were pleiotropic with phenotypes 1, 2, and 3. Phenotypes 4-13 were random in all simulations, with no genetic component. Other methods performed less well (Supplemental Table 3).

As described previously (see model score in the Method section), SAPPHO has a single tuning parameter defined by the least significant univariate p-value that can be added to an association model. Because model scores are calibrated by permutation, this parameter does not affect stringency in terms of FDR or FWER. It can affect power, however, because a more stringent threshold will reject weaker true associations. It also affects computational cost because a looser threshold yields a longer list of candidate variants. We used thresholds 5 × 10^−4^, 2.5 × 10^−4^, 10^−4^, 10^−5^, 10^−6^ (Supplementary Table 4).

Using the loose threshold of 5 × 10^−4^, SAPPHO-I outperformed MultiPhen for scenario ONE, performed equivalently for scenario TWO, and had slightly less power than MultiPhen for scenario THREE (Fig. 7). To determine whether the power disadvantage was due to the greedy search or to the statistical model, we also calculated the score of model defined by the true associations. In this case, SAPPHO performed better than MultiPhen. We conclude that SAPPHO’s performance could be improved using a more sophisticated model search, for example considering considering single and double associations in a single step, or adding backward steps.

**Figure 7:**
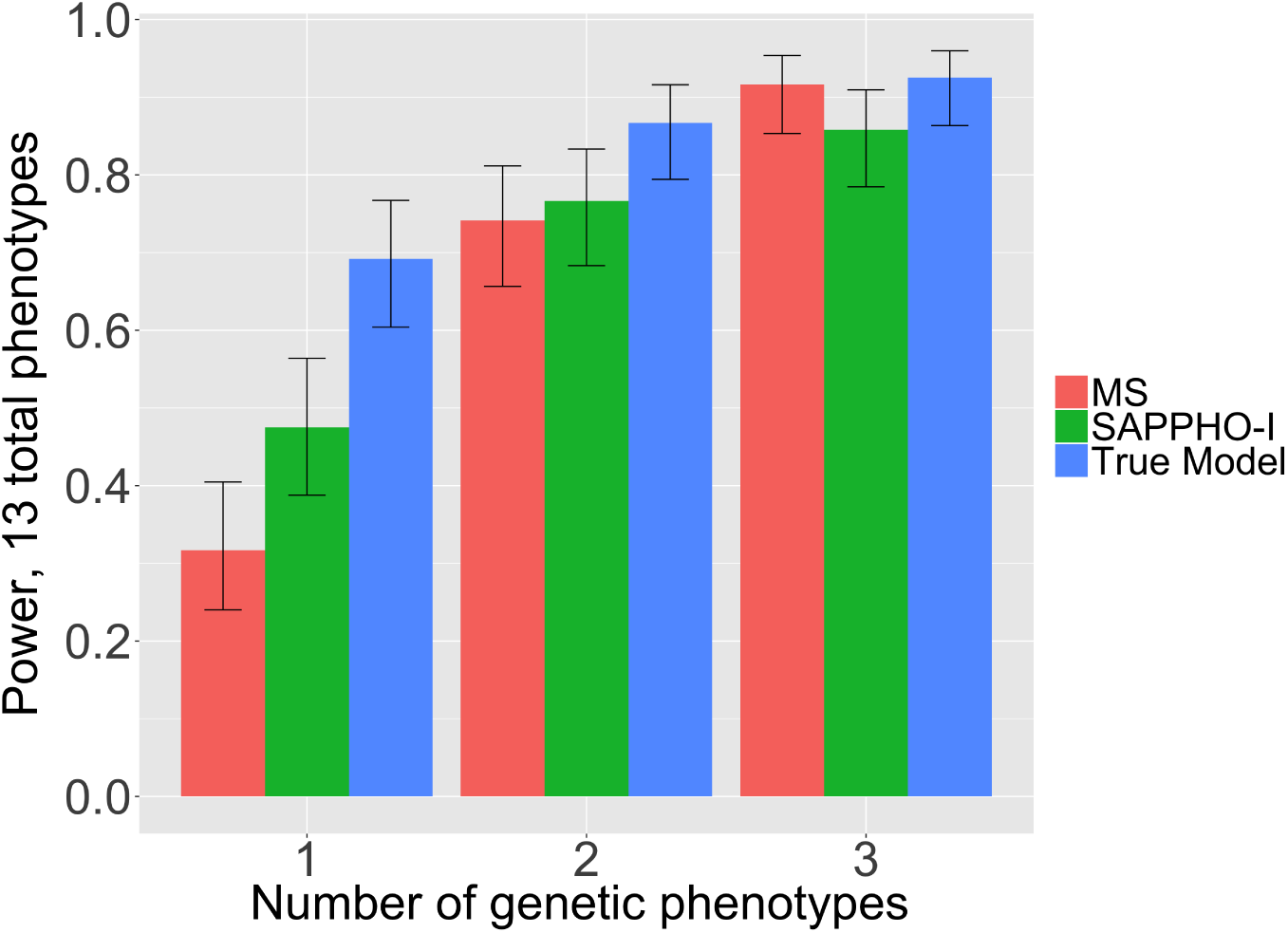
Simulations of a mixture of genetic and non-genetic phenotypes, with 1, 2, or 3 genetic phenotypes embedded as part of 13 total phenotypes. The remaining non-genetic phenotypes generated as standard normal random variables. The SAPPHO parameter was set to include associations with p-value < 5 × 10^−4^. Methods were MultiPhen in selection mode (MS), SAPPHO-I with a greedy forward (SAPPHO-I), and the SAPPHO-I score for the true model (True Model). The greedy forward search limits the power of SAPPHO-I; a more sophisticated strategy could improve its power.

We also performed simulations using the more stringent threshold of 10^−6^ for adding associations. In scenario ONE, SAPPHO still out-performed MultiPhen. In scenarios TWO and THREE, SAPPHO performed worse, in large part because the threshold prevented true associations from being considered.

At 0.05 FDR, both SAPPHO and MultiPhen returned false positive results. SAPPHO false positives tend to be spurious associations of a variant with an individual phenotype. MultiPhen tends to over-predict associations: given a true association to a phenotype, MultiPhen often predicts additional false-positive associations for the same SNP with additional phenotypes. These results support the hypothesis that the additional pleiotropic associations found by MultiPhen in the ARIC data are false positives rather than true associations, consistent with the lack of significance for these associations in the larger CHARGE data set.

## 4 Discussion

We have presented a Bayesian-motivated method, SAPPHO, designed to detect pleiotropic effects in GWAS. SAPPHO exploits previously observed association patterns to identify additional variants following the same pattern. Representative methods were selected for comparison: SHom and SHet, which pool summary statistics of all variant-phenotype associations to define a combined test-statistic; PCH, which constructs linear combinations of phenotypes for genotype data to be regressed on; and MultiPhen, which performs reverse regression such that the phenotypes are treated as predictors to explain the variance of genotypes. In applications to a real data set, ECG phenotypes from ARIC, with known positives available from the much larger CHARGE study, SAPPHO performed better than other pleiotropy methods in discovering the true associations at genome-wide significance. SAPPHO also performed best when additional random phenotypes augmented the true phenotypes, an assessment of performance when pleiotropy involves only a subset of the phenotypes in a study.

SAPPHO uses an association model that is in the exponential family, which makes it amenable to use with summary statistics rather than individual phenotype-genotype data in the context of meta-analyses. In applications to metaanalysis data from CHARGE, SAPPHO identified 295 loci at 0.05 FDR, corresponding to 171 loci in the genome-wide significance gold standard and 124 novel loci. Gene sets corresponding to cardiac electrophysiology are highly enriched for these novel loci, supporting a conclusion that SAPPHO has identified many additional relevant loci beyond those previously reported. While some of the additional loci may arise from using a less stringent 0.05 FDR threshold compared with the 0.05 FWER threshold, we also note that there are no established methods to define significant loci for pleiotropic tests. Investigating the direction of effect for pleiotropic variants, we find that variants affecting pairs of traits often have relative directions of effect that are different and that often do not match the overall phenotypic correlation. These findings suggest that multiple independent biological mechanisms connect pleiotropic variants to downstream phenotypes.

Simulated phenotype and genotype data sets were used to explore the reasons for the superior performance. These studies suggested that the version of SAPPHO that models the full phenotype covariance matrix, SAPPHO-C, can actually perform poorly when phenotypes are strongly correlated. A simplified version in which phenotypes are modeled as independent when conditioned on genetic effects, SAPPHO-I, retains robust performance. SAPPHO-I is also more computationally efficient and more amenable to use with summary data from meta-analysis.

Simulations also demonstrated that SAPPHO can have an advantage over univariate tests even applied to a mixture of phenotypes in which some lack genetic effects. The Bayesian prior learns this pattern and is able to boost associations with the phenotypes that have genetic effects.

The MultiPhen method also performed well. It out-performed SAPPHO in some simulation settings involving weaker effects, although it had a drawback of over-predicting spurious associations for variants with true associations for a subset of phenotypes.

SAPPHO depends on a single adjustable parameter, which in effect determines the minimum effect strength that can be entered into the genetic model. Permitting weaker effects, expressed as a looser univariate p-value, improved SAPPHO’s performance. We found, however, that the greedy forward search implemented by SAPPHO occasionally yields a local rather than global optimum, as assessed by calculating the score of the true model. Improving the search heuristic, for example by permitting the model to add two associations simultaneously, may improve the performance.

We conclude that SAPPHO, and particularly the SAPPHO-I implementation for summary statistics, is a powerful method for discovering pleiotropic patterns of association in the context of single studies, with access to individual genotype and phenotype data, and also to meta-analyses. Application to large compendiums of GWAS results, for example dbGaP or the UK BioBank, could lead to new discoveries of genetic associations and patterns of shared genetic architecture for human phenotypes and disease.

## Funding

This work has been supported by NSF DMS-1228248 to JSB and DEA, by NIH R01HL116747 to DEA.

## Author Contributions

JZ developed the algorithms, implemented the methods, generated the results, reviewed the results, and wrote the manuscript. The CHARGE ECG Working Group provided the gold standards and underlying data. JSB and DEA developed the algorithms, reviewed the results, and wrote the manuscript.

## Ackknowledgements

The Atherosclerosis Risk in Communities Study is carried out as a collaborative study supported by National Heart, Lung, and Blood Institute contracts (HHSN268201100005C, HHSN268201100006C, HHSN268201100007C, HHSN268201100008C, HHSN268201100009C, HHSN268201100010C, HHSN268201100011C, and HHSN268201100012C), R01HL087641, R01HL59367 and R01HL086694; National Human Genome Research Institute contract U01HG004402; and National Institutes of Health contract HHSN268200625226C. The authors thank the staff and participants of the ARIC study for their important contributions. Infrastructure was partly supported by Grant Number UL1RR025005, a component of the National Institutes of Health and NIH Roadmap for Medical Research.

The Atherosclerosis Risk in Communities study has been funded in whole or in part with Federal funds from the National Heart, Lung, and Blood Institute, National Institutes of Health, Department of Health and Human Services, under Contract nos. (HHSN268201700001I, HHSN268201700003I, HHSN268201700005I, HHSN268201700004I, HHSN2682017000021). The authors thank the staff and participants of the ARIC study for their important contributions.

The authors would also like to acknowledge NIH grant HL105756.

## Competing Financial Interests

No competing financial interests.

## Description for supplementary tables

Supplementary Table 1: Gold-standard from CHARGE meta-analysis results at different thresholds for generating the connected components.

Supplementary Table 2: SNPs detected by pleiotropy methods on ARIC data at 0.05 FWER. For each table, multiple SNPs can belong to the same locus. The number of loci detected by each method is shown in Fig. 2.

Supplementary Table 3: All variants detected by SAPPHO-I on CHARGE meta-analysis results at 0.05 FDR.

Supplementary Table 4: Results for simulation with noise phenotypes at 0.05 FDR. Sheet 1: Number of true positives and false positives for all methods; Sheet 2: Number of true positives for SAPPHO-I fed with true associations at different thresholds, and MultiPhen in select mode; Sheet 3: association patterns for SAPPHO-I and MultiPhen in select mode: *Missed entirely* denotes the number of variants missed entirely; *Underpredicted* denotes the number of variants detected but with a subset of true associations found; *Exactly-predicted* denotes variants detected with the correct association pattern; *Over-predicted* denotes variants detected with a mix of true associations and spurious associations; *False positives* denotes variants reported but lacking any true associations.

Supplementary Table 5: Gene sets enriched for loci reported by SAPPHOI at 0.05 FDR for CHARGE ECG meta-analysis, calculated for all loci and for novel findings defined by excluding the ECG gold-standard loci. Meaning of the columns: ‘Pathway’ denotes the pathway name; ‘pathway gene counts’ denotes the number of genes in that pathway; ‘SAPPHO gene counts’ denotes number of all genes detected by SAPPHO; ‘SAPPHO 11’ denotes the number of genes detected by SAPPO and in that corresponding pathway; ‘SAPPHO 10’ denotes the number of genes detected by SAPPHO but not in the pathway; ‘SAPPHO 01’ denotes the number of genes in the pathway but not detected by SAPPHO; ‘SAPPHO 00’ denotes number of genes not in the pathway and not detected by SAPPHO; ‘SAPPHO pval’ denotes the pvalue of Fisher’s exact test; ‘SAPPHO loci’ denotes the genes in the ‘SAPPHO 11’ column. The following columns are of the same meaning for the SAPPHO loci excluding the gold-standard.

